# GFETM: Genome Foundation-based Embedded Topic Model for scATAC-seq Modeling

**DOI:** 10.1101/2023.11.09.566403

**Authors:** Yimin Fan, Adrien Osakwe, Shi Han, Yu Li, Jun Ding, Yue Li

## Abstract

Single-cell Assay for Transposase-Accessible Chromatin with sequencing (scATAC-seq) has emerged as a powerful technique for investigating open chromatin landscapes at single-cell resolution. However, analyzing scATAC-seq data remain challenging due to its sparsity and noise. Genome Foundation Models (GFMs), pre-trained on massive DNA sequences, have proven effective at genome analysis. Given that open chromatin regions (OCRs) harbour salient sequence features, we hypothesize that leveraging GFMs’ sequence embeddings can improve the accuracy and generalizability of scATAC-seq modeling. Here, we introduce the Genome Foundation Embedded Topic Model (GFETM), an interpretable deep learning framework that combines GFMs with the Embedded Topic Model (ETM) for scATAC-seq data analysis. By integrating the DNA sequence embeddings extracted by a GFM from OCRs, GFETM demonstrates superior accuracy and generalizability and captures cell-state specific TF activity both with zero-shot inference and attention mechanism analysis. Finally, the topic mixtures inferred by GFETM reveal biologically meaningful epigenomic signatures of kidney diabetes.

## 1. Introduction

Recent advancements in single-cell sequencing technologies have endowed biologists with the ability to elucidate the properties of individual cells from multiple perspectives including gene expression measured by single-cell RNA-sequencing (scRNA-seq) [1] and chromatin accessibility measured by Assay for Transposase-Accessible Chromatin with sequencing (scATAC-seq) [2–5]. scATAC-seq allows us to examine the open chromatin landscapes that dictate gene regulation at the single-cell level. Effectively harnessing scATAC-seq data promises to identify cell-type-specific regulatory elements and map them to pathogenetic mechanisms. Currently, scATAC-seq is capable of profiling millions of open chromatin regions (peaks) from thousands of cells at reasonable costs. Thus, it is of vital significance to develop computational methods that can model these emerging datasets and uncover novel biological insights.

Computational modeling of scATAC-seq data is faced with many challenges. The first arises from the large number of peaks detected by statistical software based on an enrichment of aligned reads relative to background regions in the reference genome [6]. Technological advancements have led to the detection of millions of highly variable peaks in datasets containing millions of cells. This results in a high dimensional and often sparse cells-by-peaks matrix, demanding substantial computational resources. Furthermore, compared to scRNA-seq, scATAC-seq data has fewer reads per cell, and the sequenced open chromatin regions are much larger than the transcribed regions [7], leading to a lower signal-to-noise ratio [8]. Though cross-tissue and cross-species integration and transfer learning have been well explored in scRNA-seq data [9–11], it is much less so in scATAC-seq data due to the lack of a common set of features (i.e., genes used in scRNA-seq). This issue is particularly concerning given the relatively limited availability of scATAC-seq datasets. Effective transfer learning that leverages comprehensive reference scATAC-seq datasets is critical to improve analyses in under-explored diseases, tissues and species.

Several computational methods, including both statistical and deep learning methods, have been developed to analyze scATAC-seq datasets. These methods can be broadly classified into two categories, sequence-free and sequence-informed methods. The former, involving methods like PCA [12], Cicero [13], SCALE [14], and cisTopic [15], treat scATAC-seq datasets as a peak-by-cell matrix, disregarding the DNA sequences underlying each peak. Although these sequence-free methods can project cells onto a low-dimensional, biologically meaningful manifold for clustering, classification, and analysis, the omission of DNA sequence information in these methods can lead to sub-optimal performance. In contrast, sequence-informed methods like chromVAR [16], BROCKMAN [17], CellSpace [18], SIMBA [19] and scBasset [20], integrate DNA sequence information from peaks in modeling scATAC-seq datasets to encode sequence information, therefore could offer novel and more reliable biological insights. However, these methods still have major limitations. First, existing sequenced-informed methods such as scBasset [20] and CellSpace [18] cannot be applied to novel cells aside from the training set, while generalization and transfer to unseen cells is vital for developing a general computation method considering the limited sample size for some tissues and studies. Second, existing methods typically utilize k-mer or Transcription Factor (TF) motif features or employing Convolutional Neural Networks (CNN) to model the DNA sequence from peaks, which are limited to local peak sequence features and may overlook long-range sequence context. On the other hand, Genome Foundation Models (GFMs) have demonstrated promising performance on a wide range of genomics modeling tasks including predicting chromatin profiles [21–24]. We hypothesize that the expressive power brought by GFMs via large-scale pre-training can generate informative DNA sequence embeddings that innovate and improve upon existing scATAC-seq modeling approaches.

In this study, we present Genome Foundation Embedded Topic Model (GFETM), a novel and interpretable generative topic modeling framework enhanced by GFMs. Our approach builds upon the concept of an embedded topic model (ETM) [25], a powerful topic model implemented with a Variational Autoencoder (VAE). Its superior performance for single cell modeling and integration has previously been demonstrated with scRNA-seq and single-cell multi-omic data by scETM [11, 26] and moETM [27], respectively. Hence, we extend this framework to sequence-informed scATAC-seq modeling. The GFETM jointly optimizes a GFM and ETM, allowing the sequence knowledge from the GFM to be seamlessly integrated and adapted to the ETM. This integration, combined with the interpretability offered by the ETM’s linear decoder, enhances the overall performance of the GFETM. GFETM demonstrates state-of-the-art cell clustering and high scalability on large datasets due to its batch-agnostic embeddings and ability to infer cell states. It imputes peak accessibility on unseen chromosome regions and denoises raw scATAC-seq data, enhancing regulatory signals. Additionally, GFETM enables knowledge transfer across tissues, species and omics. Through zero-shot inference and post-hoc attention mechanism analysis, we validated that GFETM captures the cell-state transcription factor activities. In summary, by integrating GFMs pre-trained following the principal of LLMs, which have succeeded in may areas, we developed GFETM, which emerges as a powerful, generalized and interpretable scATAC-seq analysis framework which could facilitating a nuanced understanding of cell states and epigenomic landscapes.

## 2. Results

### Method Overview

GFETM is an interpretable and transferable framework for scATAC-seq modeling (Fig. 1). GFETM is composed of two main components, the ETM component and the GFM component, which respectively take the cell-by-peak matrix and peak-specific nucleotide sequences as input. The ETM encoder projects the peak vector of each cell onto a latent cell topic mixture through a two-layer multi-layer perceptron (MLP). The cell topic mixture is then passed to the linear ETM decoder and combined with the sequence embeddings generated by the pre-trained GFM component to compute the expected peak accessibility in each cell. The training objective is to maximize the evidence lower bound (ELBO) of the log marginal likelihood of the input cell-by-peak matrix (**Methods** 4.1). Therefore, GFETM embraces the flexibility of the non-linear encoder for cell and sequence embeddings but also retains high interpretability attributable to the linear factorization used by the decoder (**Methods** 4.3). For transfer learning as is demonstrated in Fig. 1b, the trained encoder of GFETM from one dataset can be used to project unseen cells from different species, tissues and omics onto the same topic mixture space using aligned features (**Methods** 4.4). As shown in Fig. 1c, the trained GFM and ETM can be used to impute the chromatin accessibility on unseen chromosome regions (**Methods** 4.5), denoise the original raw cell-by-peak matrix (**Methods** 4.6) as well as infer the cell-state specific TF activity (**Methods** 4.7). Furthermore, we can associate topics with cell types or conditions by differential analysis. The top peaks of the differential topics can be subject to motif enrichment to identify transcription factors and also associated with target genes to identify cell-type-specific regulatory modules (Fig. 1d).

**Figure 1:**
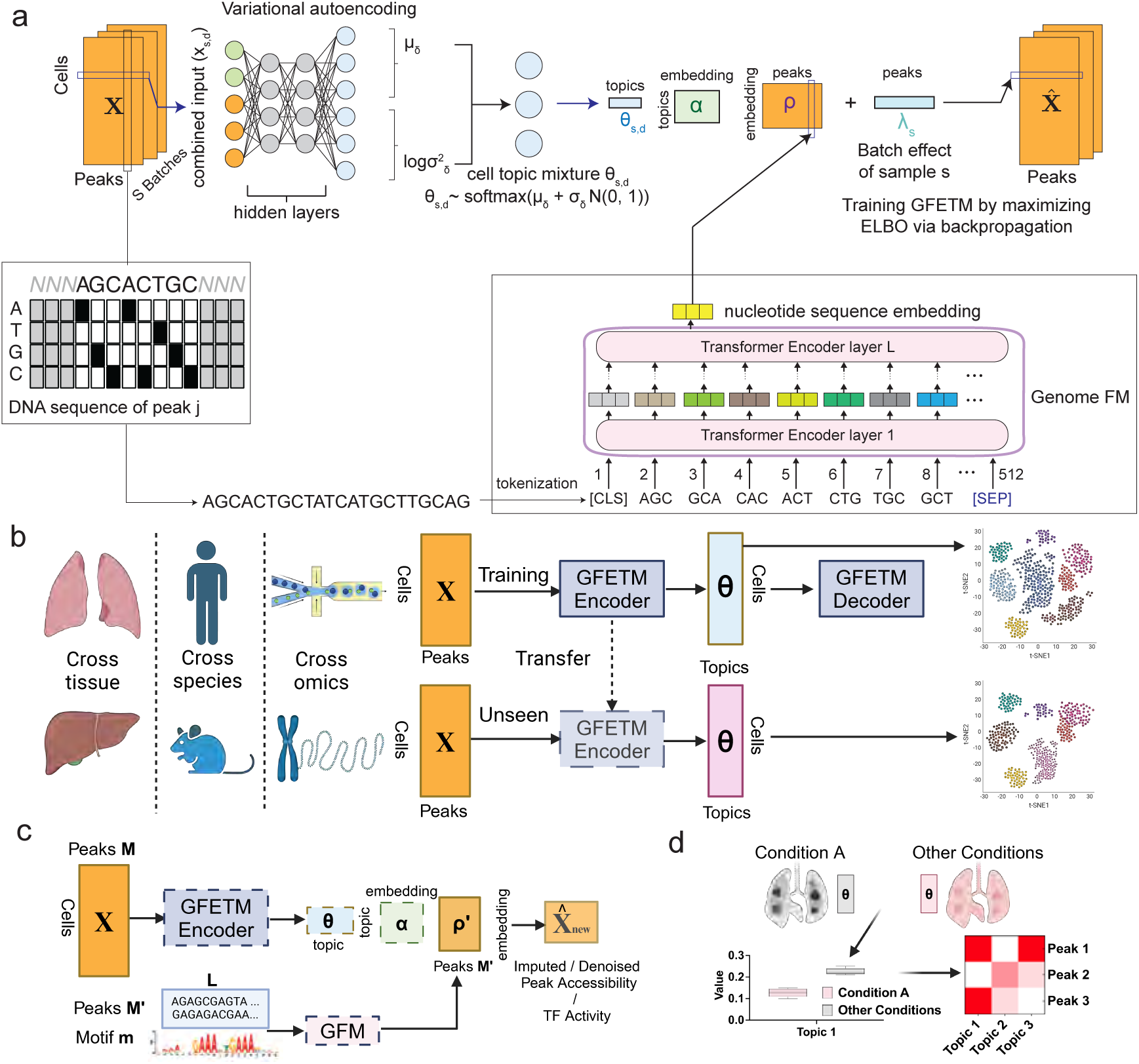
Overview of GFETM. **a.** Overview of GFETM. GFETM models the scATAC-seq data using an embedded topic-modeling approach. Each scATAC-seq profile serves as an input to a variational autoencoder (VAE) as the normalized peak count. The encoder network produces the latent topic mixture for clustering cells. The GFM takes the peak sequence as input and outputs peak embeddings ***ρ***. Given the cell topic mixture ***θ****_s,d_* of cell *d* from batch *s* and the peak sequence embedding ***ρ***, the linear decoder learns topic embedding ***α*** as its weights to reconstruct the input. The encoder, decoder and GFM are jointly optimized by maximizing the evidence lower bound (ELBO) of the scATAC data log marginal likelihood. **b.** Transfer learning of GFETM. GFETM is first trained on one tissue/species/omics and then applied to another unseen tissue/species/omics with or without further fine-tuning. c. Imputing / Denoising the peak accessibility matrix. and inferring TF activity Trained GFM and ETM can be used to denoise the original scATAC-seq matrix, impute the accessibility on unseen chromosome regions as well as infer the cell-state specific TF activity. Given the *M ^′^* sequences, we can infer their likelihood in the *N* cells by applying the dot product of the cell embedding (θ produced by the trained encoder), topic embedding (α as learned parameters), and peak sequence / motif sequence embedding (ρ*^′^* produced by the trained GFM). If *M ^′^* peaks is a subset of *M* peaks, it can be considered as denoising the original scATAC-seq matrix. Otherwise, GFETM imputes on unseen chromosome regions. If motif sequence is used as input, it can be viewed as inferring the TF activity. **d.** Differential topic analysis to identify epigenomic signatures. We first conduct differential analysis over the cell-level topic mixture θ*_k_* ∈ *R^N×^*^1^ by comparing *N ^′^* cells from one cell type or condition against the rest of the *N* − *N ^′^* cells. We then analyze the *K^′^* differential topic distributions (***β*** ∝ ***αρ***) by identifying the top peaks that exhibit the highest topic scores, which can be linked to relevant TFs or genes.

### Evaluating sequence embeddings from diverse GFMs

We first evaluated ETM using the nucleotide sequence embeddings from different GFMs to determine if embeddings from pre-trained GFMs contain useful information for scATAC-seq analysis. To comprehensively compare GFMs with different scales and sizes and understand their impact on scATAC modeling, we harnessed 13 GFMs and compared their performance under two settings: 1) Fixing the peak embeddings ***ρ*** to the sequence embeddings from GFMs during ETM training; 2) Initializing the peak embeddings ***ρ*** with the sequence embeddings from GFMs and updating them during the ETM training. The results on three common benchmarking datasets are depicted in Fig. 2. We used Adjusted Rand Index (ARI) as a metric to benchmark unsupervised cell clustering. Compared to random initialization, initializing with peak sequence embeddings from GFMs consistently led to higher ARI when the peak embeddings were frozen during training. This indicates that leveraging the latent information from GFM embeddings benefits cell clustering when compared to random embeddings. Moreover, as the dimension of the peak embeddings increases, the models’ performance improves due to the higher expressiveness of the cell representation.

**Figure 2:**
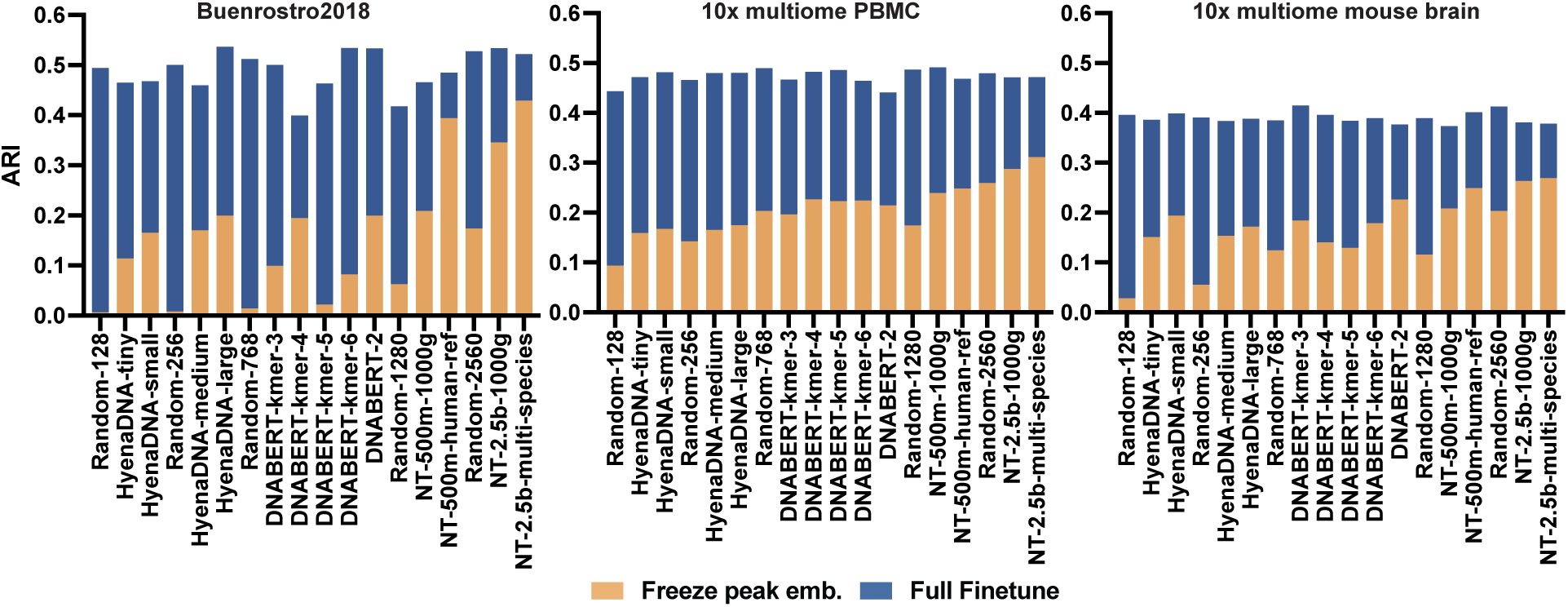
Cell clustering evaluation of GFETM using peak embeddings from different GFMs. We evaluated the performance of using frozen peak embedding (beige bars) and fine-tuning peak embeddings during GFETM training (blue bars). For comparison of model performance as a function of peak embedding size, we ordered from left to right the GFMs by peak embedding size after the random peak embedding. For example, both HyenaDNA-tiny and HyenaDNA-small have peak embedding size of 128, and HyenaDNA-medium has embedding size of 256, and so on (where ‘tiny’, ‘small’, and ‘medium’ indicate the pre-training sequence lengths of 1k, 32k, and 160k, respectively.). The largest embedding size is 2560 corresponding to the last 3 bars. Adjusted Rand Index (ARI) on the y-axis was used to evaluate the clustering by the Louvain algorithm based on the corresponding cell embeddings from each framework for 3 separate datasets as shown in the 3 panels.

Additionally, a noteworthy observation is the alignment between the performance of GFMs and cell clustering outcomes. For instance, NT-2.5b-multi-species performs exceptionally well across all released Nucleotide Transformers [23] checkpoints, and also demonstrates superior performance when its embeddings are used with an ETM for cell clustering. NT-500M-human-ref outperforms NT-500M-1000g by a large margin, especially on Human HSC Differentiation dataset, which may be the result of better alignment between the peak sequences (hg38/hg19 reference genome) and the pre-training reference genome (hg38 reference genome) than with the 3202 individual genomes from the 1000 Genome Projects.

However, directly utilizing the frozen peak embeddings from GFMs does not yield significant improvements in model performance over an ETM that optimizes randomly initialized peak embeddings (Fig. 2). This suggests that the information from GFMs is not comprehensively utilized by the ETM. To address these drawbacks, GFETM jointly and efficiently trains the GFM and ETM components, thereby fully integrating sequence knowledge from pre-trained GFMs (**Methods** 4.3). Indeed, after fully fine-tuning the GFM and ETM weights end-to-end, we observe drastic improvements (Fig. 3a). In the following experiments, we chose the pre-trained DNABERT [21] as the GFM component in GFETM as its light-weight architecture reduces computation and is based on a standard transformer.

**Figure 3:**
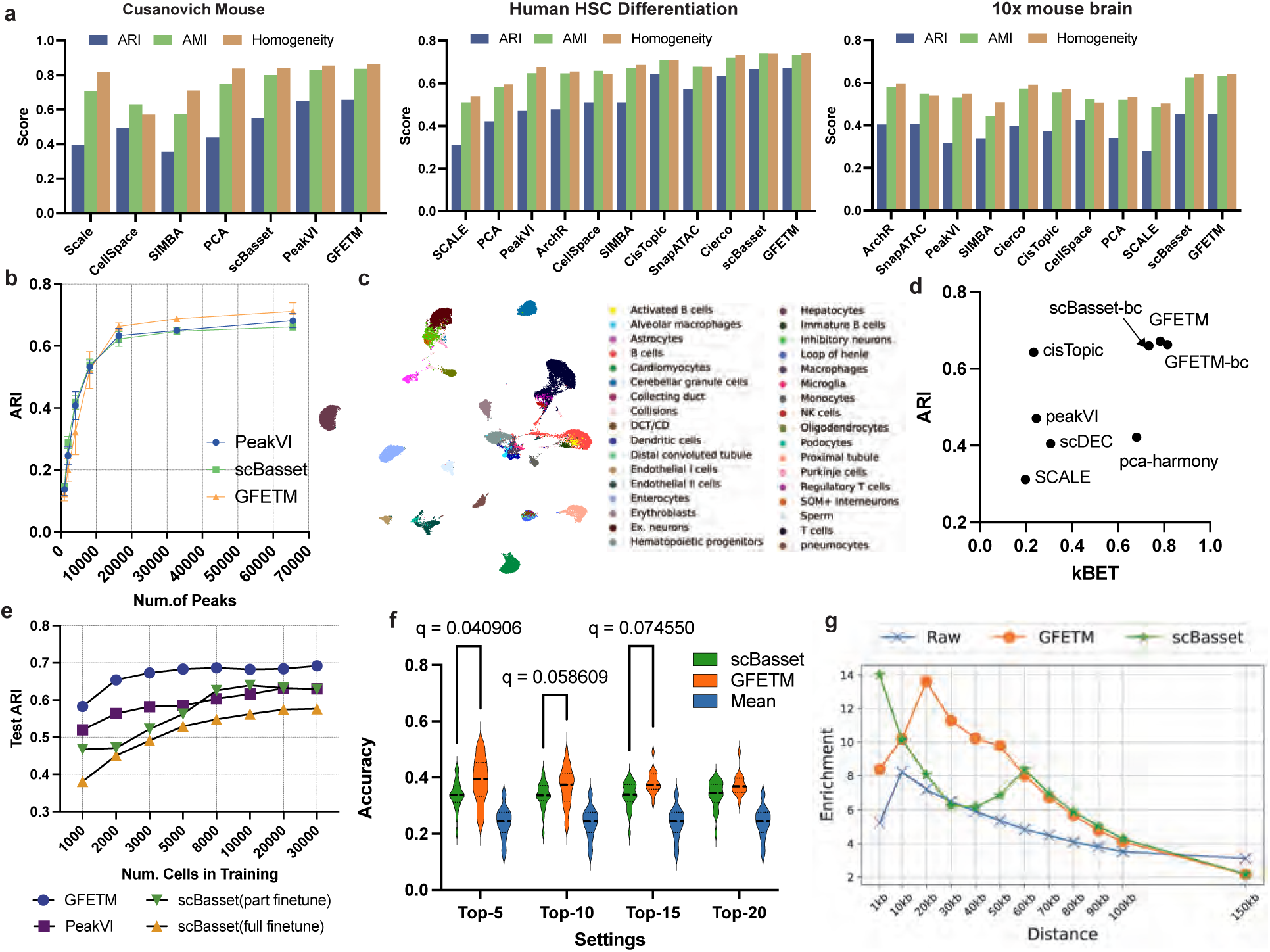
Benchmarking GFETM against existing methods. **a.** Performance on Human HSC Differentiation, 10x mouse brain and Cusanovich Mouse. ARI, AMI and Homogeneity were used as quantitative evaluation metrics. **b.** Scalability on large-scale datasets. The average results for 20000 cells, 40000 cells, 60000 cells and 80000 cells from the Cusanovich-Mouse dataset. scBasset and PeakVI are used as baseline methods. **c.** Visualization for large-scale integration of GFETM. **d.** Biological conservation and batch effect correction. For each method, we computed the ARI as the biological conservation and kBET as the batch effect correction metric. Both scores are the higher the better. scBasset-bc denotes the batch correction version of scBasset. GFETM-bc fits the linear correction factor *λ*. **e.** Generalization on unseen cells during the training. ARI on 10K unseen cells cells as a function of the number of training cells, ranging from 1K to 30K cells. Both the training and test set were obtained from the Human HSC Differentiation dataset. **f.** Imputing the accessibility on unseen chromosome regions. We performed the experiments in a leave-one-out cross-chromosome imputation setting. The results were averaged over 22 chromosomes. Top-*k* accuracy is defined as the proportion of true positives in the top-*k* imputed accessible peaks. As a negative control, we also evaluated mean prediction (the average frequency of the test peaks on cells). **g.** Denoising the raw scATAC-seq matrix for HSC cells. We first detected marker peaks for specific cell type using raw, GFETM, or scBasset denoised scATAC signals. We then computed the marker gene enrichment based on the genes whose TSS is within a defined genomic distance of the marker peak. x-axis is the distance threshold of marker peaks to the TSS of marker genes.

### GFETM improves scATAC-seq cell representation

To validate the effectiveness of GFETM in learning meaningful cell representation, we first assessed the performance of GFETM on three standard benchmarking datasets [7] and compared its performance against other baseline methods (Fig. 3a; **Methods** 4.14). We evaluated the quality of the inferred cell embeddings by comparing Louvain clustering results with ground-truth cell-type labels using the ARI, Adjusted Mutual Information (AMI) and cell type Average Silhouette Width (ASW). Sequence-informed deep learning methods, including our GFETM and an existing state-of-the-art method, scBasset, outperform all other baseline methods by a large margin, which demonstrates the advantage of integrating peak DNA sequence information into scATAC-seq modeling. GFETM achieves competitive performance when compared with scBasset in the majority of tasks. We also observed substantial relative improvement of GFETM over most baseline methods (Supplementary Fig. S1). In addition, we also experimented with an ETM model using a CNN architecture (Supplementary Section S1.1; Supplementary Fig. S2) instead of the GFM and observed inferior performance (Supplementary Fig. S3). We also evaluated the impact of hyper-parameter tuning on cell clustering performance and showed that GFETM is robust to different hyper-parameter settings (Supplementary Section S1.4 and Supplementary Fig S4).

Furthermore, several scATAC-seq benchmarking datasets were limited in the number of cells profiled (e.g. less than 10k) and sequencing depth [7], while the recent advancements in high-throughput sequencing technologies enables the generation of much larger datasets. To evaluate the scalability of GFETM on large-scale datasets, we utilized the Cusanovich-Mouse dataset. We compared the performance of GFETM with the state-of-the-art (SOTA) methods scBasset and PeakVI (Fig. 3b). We varied the number of cells in the dataset from 20k to 80k and adjusted the number of selected highly variable peaks from 1k to 70k and showed the average results across cell numbers. Results with respect to each cell number are shown in Supplementary Fig. S5. GFETM consistently outperformed scBasset and PeakVI across all datasets, and performed particularly well in scenarios with a large number of peaks, highlighting its capability in fully utilizing the DNA sequence information from OCR with GFM. We evaluated the robustness of results with respect to clustering hyperparameters in Supplementary Table S5 and S6.

Additionally, we visualized the UMAP of cell embeddings from GFETM on the Cusanovich-Mouse dataset (Fig. 3c), which qualitatively illustrates GFETM’s ability to reliably stratify cell types. We confirmed the reported performance by visualizing the clustering across the entire atlas by tissues and Louvain clusters in Supplementary Fig. S6. We also visualized the cell embeddings on the Cusanovich-Mouse dataset for baseline methods in Supplementary Fig. S7 and Supplementary Fig. S8. We then evaluated the runtime and memory usage of GFETM on large-scale datasets in Supplementary Fig. S9. We found that the runtime and memory usage of GFETM is reasonable even when applied to large scale datasets. The detailed analysis is shown in Supplementary Material S1.2.

A single-cell ATAC-seq dataset may contain cells from multiple batches, donors and/or experiments. To enable accurate cell modeling, it is important to strike a balance between batch effect removal and the conservation of biological variation. To this end, we evaluated the biological variation conservation (ARI) and batch effect removal (kBET) performance of existing models on the Human HSC Differentiation dataset (Fig. 3d). We found that GFETM achieves the highest ARI and the second highest kBET even without including the batch term *λ* in the model (**Methods**), which indicates that GFETM can generate high-quality batch-agnostic cell embeddings. After adding the batch removal term *λ* in the model (Eq 4), GFETM-bc, we achieve the best performance on kBET with a slightly lower ARI. Therefore, both GFETM and GFETM-bc are competitive in cell type stratification and batch correction (Fig. 3d; Supplementary Fig. S10). As GFETM already strikes a good balance between batch correction and biological diversity conservation without the batch effect term, we chose to omit the term for subsequent analyses.

Generalizability is an important concern in scATAC-seq cell representation methods, as generalizable models could be directly applied to newly sequenced cells without the need to retrain the model. Some existing methods such as CellSpace and SIMBA cannot be generalized to unseen cells directly. We compared the generalizability of our method with scBasset and PeakVI on unseen cells. Both the training cells and test cells came from the Cusanovich-Mouse dataset. The test set contains 10k cells. Since scBasset treats the cell embeddings as the model parameters as opposed to amortizing them on the encoder (i.e., our VAE-based GFETM and PeakVI frameworks), it is not directly transferable. As a result, we needed to fine-tune scBasset both partly and fully on the test set. We employed two strategies to adapt scBasset on unseen cells. As scBasset contains a peak encoder and a peak classifier, the first strategy is to finetune both of them on the test set (i.e., full finetune). The second strategy was to freeze the peak encoder trained on the training set and only update the peak classifier on the test set (part finetune). We directly applied GFETM and PeakVI to the test set. GFETM demonstrated the best generalization when trained on datasets with 1k-30k cells and outperformed the scBasset models that were fine-tuned and fully trained on the test dataset (Fig. 3e).

scATAC-seq datasets are inherently noisy due to the low read depth, technical variability and biological stochasticity, thus imputation and denoising are essential to enhance the robustness of signal detection and downstream analysis. First, we evaluated the generalization of chromatin accessibility prediction on unseen peak sequences to simulate the scenario with peak sequence dropouts. Concretely, given the GFETM trained on a scATAC-seq dataset using *N* cells and *M* peaks from all but one chromosome, we assessed the accuracy of inferring the *M ^′^* unseen peak accessibility on the held out chromosome for each individual cell, which is a challenging out-of-distribution setting. The implementation details are described in **Methods** 4.5 and Supplementary Fig. S11. We performed evaluation using leave-one-chromosome-out cross validation on the Buenrostro2018 dataset. We kept peaks from the 22 autosomes and omitted chromosome X and Y. We used peaks from 21 chromosomes for training and the remaining chromosome for testing. The top 500 highly variable peaks for each chromosome were used to balance the dataset. We used the top *K* accuracy (also known as top *K* precision) as the evaluation metric. For each cell, we selected the top *K* peaks from the *M ^′^* peaks and computed the proportion of true positives among the top *K*. We then averaged the top *K* accuracy across all of the *N* cells. This metric is suitable for downstream experimental validation as only a limited number of peaks can be validated making it crucial to have high precision among the top *K* most confident predictions. We only compared the performance of GFETM with scBasset, as CellSpace, SIMBA and PeakVI cannot perform this task. We experimented prediction accuracy for *K* = 5, 10, 15, and 20 (Fig. 3f and Supplementary Fig. S12). While predictions tends to vary from chromosome to chromosome, GFETM generally outperforms scBasset by a large margin (10% for *K* = 5, 6% for *K* = 10 and 3% for *K* = 20). As a baseline, we also evaluated the average frequency of the test peaks among the test cells, confirming that both scBasset and GFETM made sensible predictions.

Besides the quantitative evaluation on imputing unseen peak chromatin accessibility accuracy, we further evaluated GFETM in denoising and enhancing the biological signal detection from scATAC-seq dataset. Concretely, we evaluated the capabilities of GFETM in denoising raw scATAC-seq counts and enhancing the signal from cell-type specific marker genes in the Human HSC Differentiation dataset. We chose HSC (Hematopoietic stem cells) cells for this analysis as it contains the most annotated list of marker genes in the dataset. We performed differential analysis on the raw and denoised counts and identified the differentially accessible peaks (i.e., marker peaks) (FDR *>* 0.1 and log-fold changes *>* 0 for Fig. 3g). We then computed the enrichment of the differential peaks with respect to marker genes (peaks within distance from 1kb to 150kb upstream of TSS region). Details are described in **Methods** 4.6. GFETM-denoised scATAC-seq signals increased the marker genes enrichment by a significant margin compared to the raw signals. We found that scBasset achieved higher enrichment for marker genes within 1 kb distance from the TSS but lower enrichment compared to GFETM beyond 1 kb. This may imply that scBasset tends to capture local promoter-proximal motif information, which can be attributed to the local receptive field of the CNN. In contrast, GFETM results were enriched for peaks at *>* 10 kb, which corresponds to distal enhancers. This could be attributed to the attention mechanism used in the GFM component, which is capable of capturing complex regulatory elements within peak regions beyond the CNN local filters. Enrichment scores under different hyper-parameter settings are shown in Supplementary Fig. S13. We found that GFETM consistently achieved better performance, especially for peaks that were distant from the TSS. Results for other cell types with annotated marker genes are shown in Supplementary Fig. S14 and demonstrate GFETM’s superiority across different cell types.

### Transferability across tissues, species, and omics

Given the superior performance of GFETM for batch correction and generalization, we conducted further experiments to assess the transferability of GFETM on datasets involving different tissues, species, and data modalities (i.e., transferring between scRNA-seq and scATAC-seq data), both in zero-shot and continual training settings. The capabilities to generalize across tissues and species is essential for improving the embedding quality especially when the training data is scarce for some tissues or species. Despite various efforts in scRNA-seq analysis [11, 28, 29]), few studies comprehensively performed cross-species and cross-tissue transfer learning using scATAC-seq datasets [30].

First, We performed zero-shot transfer learning across mouse (Fig. 4a) and human tissues (Supplementary Fig. S15), respectively. GFETM demonstrated superior zero-shot transfer learning performance compared to PeakVI (Supplementary Table S2 and S3). As expected, transfer learning onto target tissues with similar functions as the source tissues led to better results compared to distinct tissues (Fig. 4a, Supplementary Fig. S15). For example, the Pre-frontal Cortex, Cerebellum and Whole Brain all contain cells in the nervous system, whereas Bone Marrow, Spleen and Lungs are all closely related to the immune and hematopoietic systems; the liver and kidney are both key organs for the regulation of homeostasis and metabolism in the body, and are connected via the ornithine cycle. These tissues contain similar cell types and thus contain more transferable information. We used ARI to verify the positive correlation between cell-type similarity and zero-shot transfer-learning performance. (Fig. 4b; **Methods** 4.11). In the Transfer+Train setting, for each target tissue, we selected the most similar tissue as the source tissue and observed improved performance over training directly on the target tissues in the majority of the cases (Supplementary Fig. S16). We then visualized the Transfer+Train clustering (Fig. 4c). We found that the Transfer+Train from Whole Brain to Cerebellum reliably separates Cerebellar granule cells, inhibitory neurons and Astrocytes due to both tissues being primarily composed of these cell types. For the Transfer+Train between Bone Marrow and Spleen, the cell cluster containing Erythroblasts and Hematopoletic progenitors are more distant from other clusters, which can be attributed to the high prevalence of these cells types in Bone Marrow. The cell type composition of each tissue is shown in Supplementary Fig. S17.

**Figure 4:**
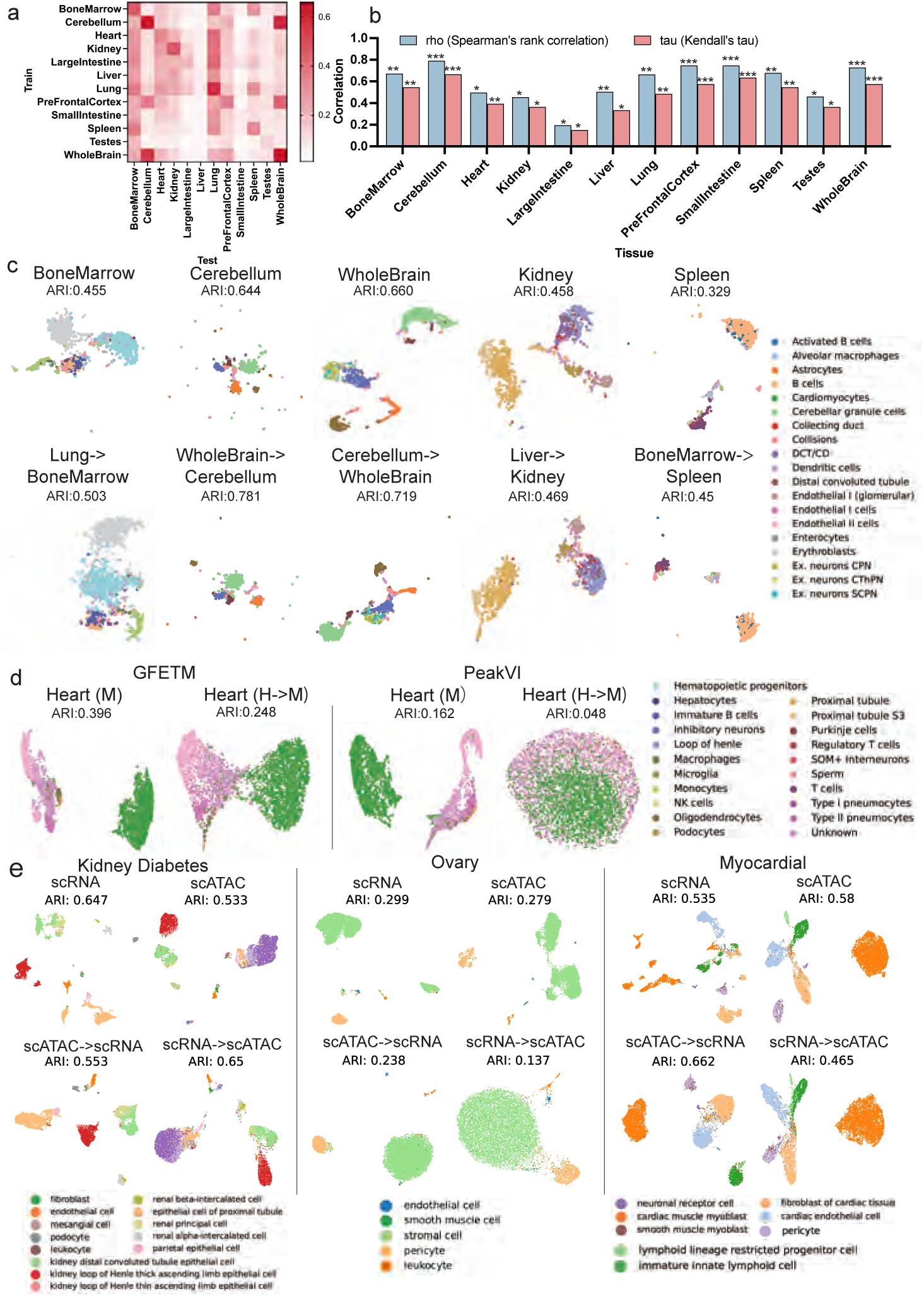
Knowledge transfer capabilities of GFETM. The performance are measure by clustering ARI. **a.** Zero-shot transfer learning on mouse tissues: GFETM trained on one tissue and tested on another. Darker heatmap colors indicate better performance. **b.** Correlation between cell type similarity and transfer performance across mouse tissues. Significance is shown by asterisks (* p=0.1, ** p=0.001, *** p=0.0001). **c.** Cross-tissue transfer: GFETM trained on one tissue, then fine-tuned on another, and tested on held-out cells. **d.** Cross-species transfer between human and mouse tissues for heart, lung, and intestine, comparing ARI and UMAP clustering with PeakVI. **e.** Cross-omic transfer on kidney, ovary, and myocardial data. UMAPs show within- and between-omic clusterings.

We then conducted zero-shot transfer learning between human and mouse tissues. Human tissues were used as the source, while mouse tissues served as the target. As shown in Fig. 4d and Supplementary Fig. S18), GFETM successfully delineated major cell types with a high ARI, achieving around 5-6 times increase compared with baseline method. For instance, in the transfer learning task involving mouse heart tissue, the GFETM trained on human heart tissue generated high-quality cell embeddings that clearly separated major heart cell types such as Cardiomyocytes, Endothelial I cells, and Endothelial II cells (Supplementary Fig. S19). Therefore, our results show that GFETM can reliably transfer single cell chromatin accessibility between human and mouse samples despite evolutionary differences.

Furthermore, we performed zero-shot cross-omic transfer learning between scRNA-seq and scATAC-seq datasets on the same tissues and species. We first aligned the scATAC-seq datasets and scRNA-seq datasets on a common set of gene features (**Methods** 4.4). GFETM trained on one modality can generate reasonable cell clusters on the other modality (Fig. 4e). For example, on an Ovary dataset, GFETM trained on scATAC-seq datasets clearly separated stromal cells and pericyte cells in scRNA-seq datasets, but did not reliably stratify rarer cell types. Interestingly, on the myocardial datasets, transfer learning from scATAC-seq datasets on scRNA-seq data yielded better clustering performance compared with directly training on scRNA-seq datasets. Indeed, myocardial scRNA datasets have large batch effects with respect to the age group of the donor, while myocardial scATAC data were less affected (Supplementary Fig. S20).

We then examined whether the clustering is affected by donors. To mitigate the impact of age, we chose donors in the same age group (44-year-old and 63-year-old for the Myocardial dataset and 62-year-old for the Ovary dataset). The 44-year-old group in the Myocardial dataset contains 3 donors and the 63-year-old group contains 2 donors. Comparing the UMAP for no-transfer against with-transfer (Supplementary Fig. S21a versus b) for the same age group of 63-year-old, the cross-omic transfer setting is less affected by the batch effects. For example, in the no-transfer setting, immature innate lymphoid cells form two separate clusters for two different donors, while in the cross-omic transfer setting, immature innate lymhoid cells form one distinct cluster with cells from both donors mixing together. Similarly, for the 44-year-old group, the cardiac muscle myoblast cells are less affected by batch effects in the cross-omic transfer setting (Supplementary Fig. S21c,d). Similar observations were made in the ovary dataset (Supplementary Fig. S22). This suggests that cross-omic transfer learning mitigates the donor-specific batch effects within the same age group. Therefore, we posit that scATAC-seq datasets harbor extensive information crucial for differentiating cell types, enabling precise clustering when applied to scRNA-seq datasets. On the other hand, we admit that our analysis was not systematic but rather illustrative, based on several example datasets. Systematically linking cell-type-specific regulatory elements to target genes [5] is required to derive further insights into the intrinsic regulatory programs.

### GFETM captures cell-state specific TF activity

GFETM encodes peak DNA sequences using GFM, enabling it to capture sequence information related to TF binding. To test whether GFETM can capture cell-state specific TF activity, we conducted two types of analyses: zero-shot inference and post-hoc attention mechanism analysis. Several existing methods were proposed to infer the TF activity from scATAC-seq datasets [31].

Our GFETM can directly use its GFM to generate motif sequence embeddings and scores TF activity (Fig. 5a). We use a multiome dataset [32] to assess the predicted TF activities using the corresponding TF expression measured in the same cell. As shown in Fig. 5b, GFETM outperforms baseline methods, including scBasset and CellSpace, in terms of correlation with TF expression in scRNA-seq datasets. For example, the TF gene MEF2C is highly expressed in GluN4 and GluN5 cell type (5c), which is consistent to our predicted inferred TF activity. In contrast, scBasset predicts high TF activity for EC/Peric and nIPC/GlN1 cells, which do not match the observed gene expression. Additional examples of other TFs are provided in Supplementary Fig. S23, S24 and S25.

**Figure 5:**
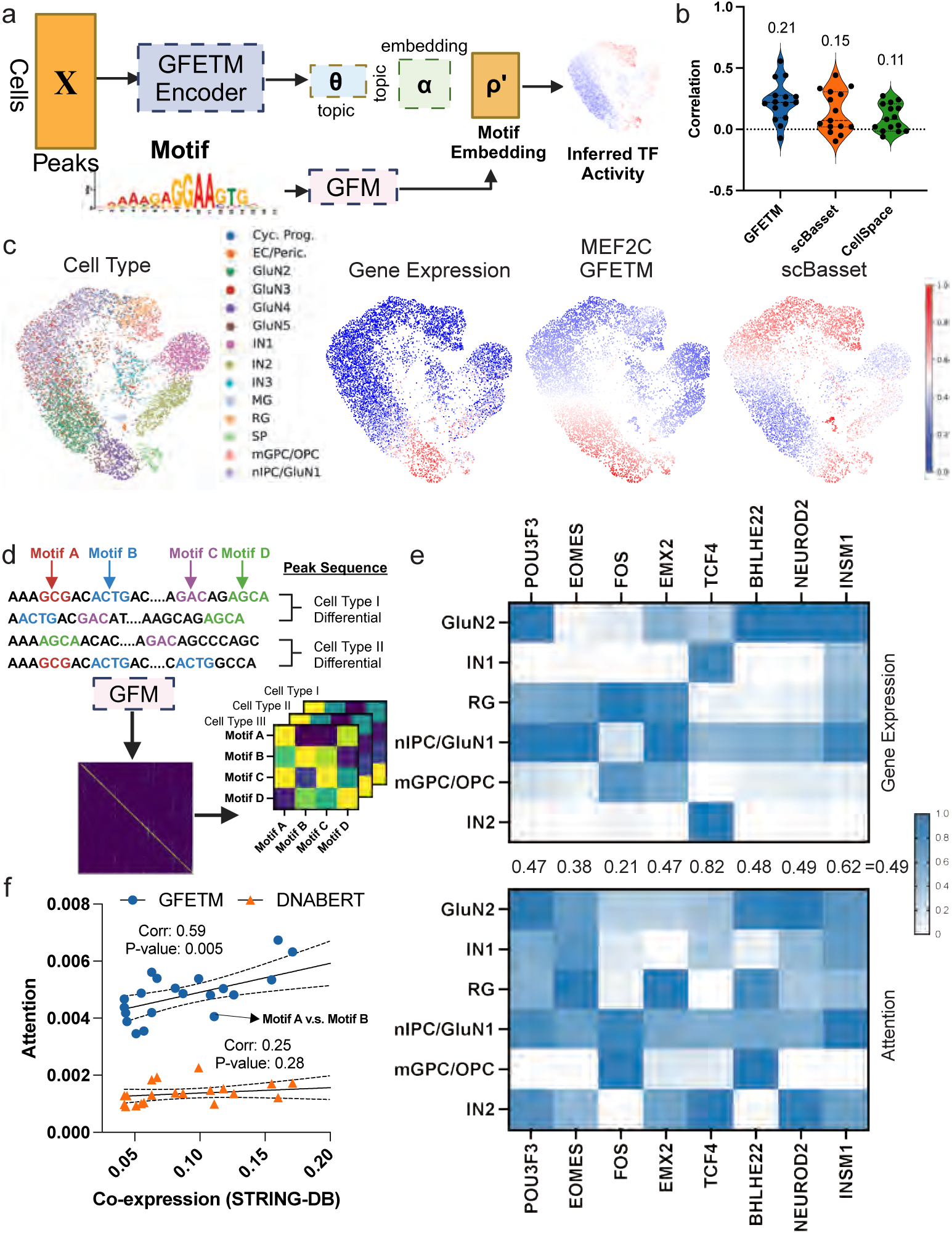
GFETM captures cell-state specific TF activity. **a**. Illustration of using GFETM for scoring the motif activities. **b**. The correlation between motif scores and gene expression of GFETM and baseline methods on signature TF motifs of human cortex. **c**. UMAP Visualization of the motif scoring result for MEF2C TF. **d**. Illustration of the interpretation of GFETM attention mechanism. Cell-type-specific attention maps were first extracted from the marker peaks using the trained GFM component and aggregated based on motif regions. **e**. The gene expression matrix (top) and attention weight matrix (bottom) with respect to six cell types in human cortex. The numbers indicate the correlation between gene expression and attention weight for each TF. **f**. The correlation between the co-expression weight from STRING database and attention weight from DNABERT and GFETM. Each point indicates a motif pair.

The attention mechanism within the transformer blocks of GFM helps capturing the connections between gene regulatory elements in the same chromatin open region. To this end, we investigated whether the attention weight on TF motif regions could reflect the TF activities. We extracted the attention maps from the last layer of the fine-tuned GFM component from GFETM for the cell-type-specific marker peak region in the scATAC-seq dataset and aggregated them by the known motif regions (Fig. 5d; Section 4.8). The marker peaks were each cell type for determined against the rest of the cell types (Wilcoxon test, *p*-value < 0.05; log-fold-change > 1.5; Supplementary Fig. S26a). We analyzed whether the attention weights for certain TF regions in cell-type specific peaks are correlated with the cell-type specific TF gene expression. We visualized the cell-type specific gene expression (top) and attention weights (bottom) of six cell types with the most differential peaks in Fig. 5e. We observe a strong correlation pattern between the two. The attention weight of GFETM has substantial cell-type specificity, similar to the gene ex-pression counterparts. For example, POU3F3 is highly expressed and has high attention weight in GluN2 and nlPC/GluN1 cells. TCF4 has high expression and attention weight in interneuron cells (IN1 and IN2). Interestingly, attention patterns may reveal transcription regulation signals not apparent from gene expression. For example, NeuroD2 is not highly expressed in IN2 (interneuron) cells compared with other cell types but with high attention weight. The regulatory role of NeuroD2 in promoting the survival and terminal differentiation has been documented [33]. Similarly, BHLHE22, a bHLH transcription factor, plays an important role in controlling OPC differentiation [34]. A baseline visualizing the correlation between the gene expression and attention weights from DNABERT is shown in Supplementary Fig. S26d.

In addition, we explored whether the attention weights between TF regions could reflect the co-expression patterns of TF genes. We collected gene co-expression patterns from STRING database [35]. We observe prominent correlation between the co-expression scores from STRING database and attention weight (Pearson correlation 0.59; Chi-squared test p-value 0.005; Fig. 5f). After training on the domain-specific human scATAC-seq dataset [32], the attention pattern from GFETM shows a more significant correlation with STRING database co-expression scores, highlighting its capability to capture intrinsic dependencies and relationships in TF motif regions within peak sequences. We also validated these findings using the co-expression patterns obtained from the paired scRNA-seq modality [32] and observed consistent improvement over the baseline DNABERT (Supplementary Fig. S26b and c).

### GFETM topics reveal epigenomic signatures of kidney diabetes

We analyzed GFETM topics that are condition-specific or cell-type specific in a kidney diabetic dataset [36]. To annotate each topic, we conducted two analyses. First, we visualized the topic distribution of 10,000 randomly sampled cells (Fig. 6a). Cells with different cell types and conditions exhibit distinct topic intensities. Second, to quantitatively identify cell-type-specific topics, we conduct differential expression (DE) analysis comparing the topic scores between cells of one cell type or condition against the rest of the cells (Fig. 6c; **Methods** 4.9). For the cell-type-specific or disease-specific topics, we identified their signature genes based on the in-cis genes of the top peaks (Fig. 6b). For instance, topic 9 is the most up-regulated in fibroblast cells and its top peaks are aligned with genes *CRISPLD2*, *ADAMTS2*, and *FBLN5*, which are supported by the literature: *CRISPLD2* plays an important role in the expansion, migration, and mesenchymal-epithelial signaling of human fetal lung fibroblast cells [37]; *ADAMTS2* is highly expressed in fibroblasts [38]; *FBLN5* is strongly associated with fibrosis disease and fibroblasts [39]. Indeed, by the same DE analysis, we observed that topic 9 is also significantly associated with kidney diabetes (Fig. 6c). As another example, topic 10 is associated with podocytes, which are highly specialized epithelial cells that cover the outside of the glomerular capillary. Its top regions are aligned with genes *PTPRO*, *LMX1B*, and *FGF1*. Previous studies indicate that abnormal expression of *PTPRO* is related to the extensive microvillus transformation of podocytes [40]. *LMX1B* is essential for maintaining differentiated podocytes in adult kidneys [41]. *FGF1* is associated with several pathways related to podocytes [42]. Moreover, we identified topic 0, 29, 38, and 55 as DE and up-regulated topics for kidney diabetes (Fig. 6c). As the most significant topic, topic 38 is endowed with top regions aligned with *PRKCQ*, *FYB1*, and *ITK*. PRKCQ has been shown to regulate kidney cancer [43], while ITK plays an important role in the up-regulation of glomeruli [44]. The biological processes involving the *FYB1* and kidney disease may deserve further investigation.

**Figure 6:**
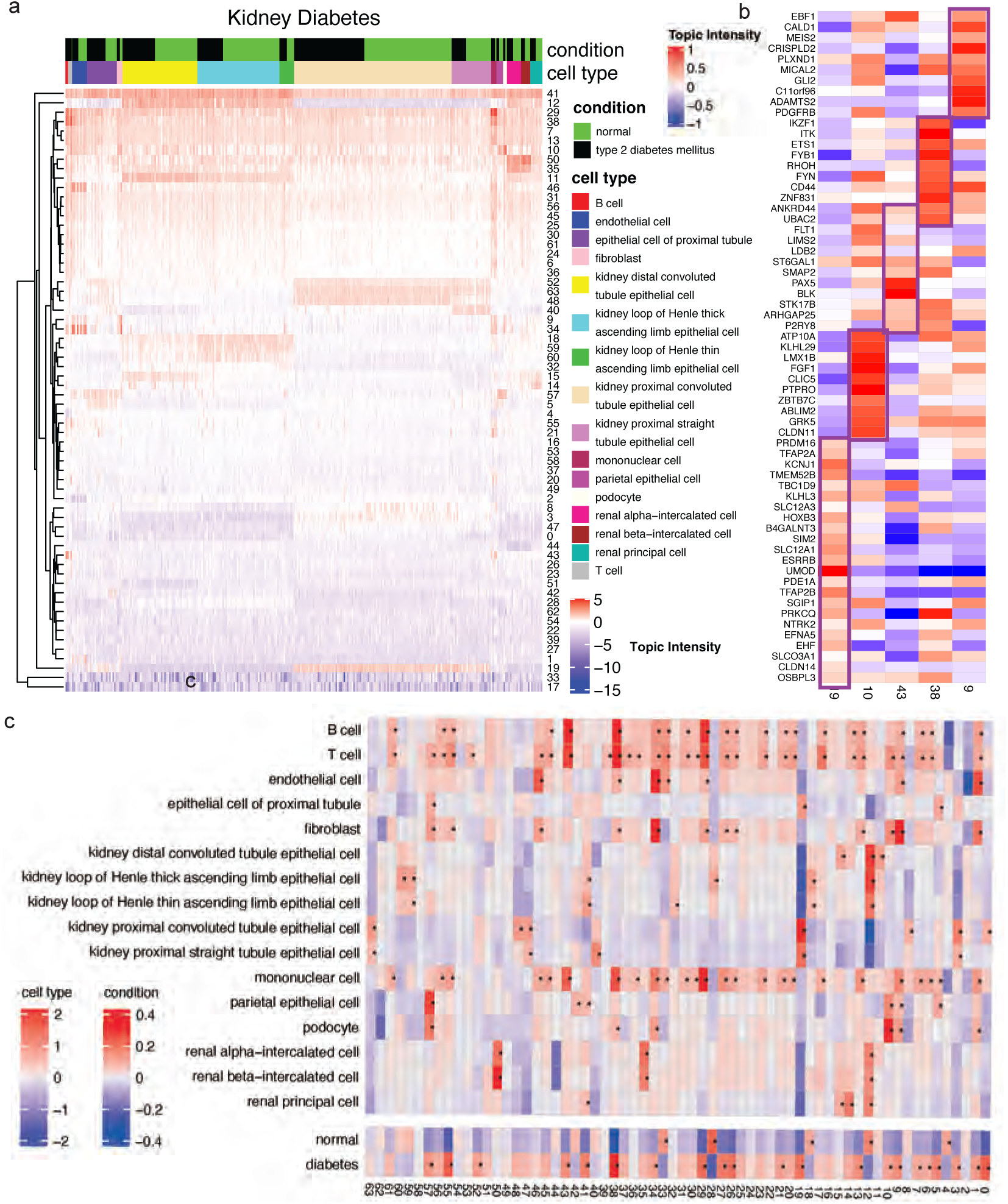
Epigenomic signatures of kidney diabetic discovered from GFETM. **a.** Topic distributions of randomly sampled 10,000 cells. Columns are cells and rows are topics. The top color bars indicate the phenotype condition and cell types. **b.** Genes aligned with the top peaks for selected topics. The aligned genes of the top peaks under 6 selected topics are shown in heatmap, where the intensity is proportional to the corresponding standardized topic scores (i.e., *β̃_k,p_* = ***α****_k,._****ρ****_.,p_*). **c.** Differential analysis to detect cell-type-specific and disease-specific topics. For each cell type or condition, we compared the topic scores of cells from the cell type or condition against the topic scores of the rest of the cells. To assess the statistical significance of the difference, we performed 100,000 permutations to construct an empirical null distribution. The asterisks indicate permutation test q-values *<* 0.1 after Bonferroni correction for testing all cell types over all topics.

## 3. Discussion

As scATAC-seq technologies mature, large-scale, high-quality datasets are beginning to be released for multiple species, tissues and conditions. This calls for the development of powerful, transferable, and interpretable representation learning methods to elucidate regulatory mechanisms from the vast amount of scATAC-seq data. To address these challenges, we developed GFETM. Compared to existing methods, our main contribution is the use of a pre-trained GFM to learn sequence embeddings that helps model scATAC-seq data. To the best of our knowledge, GFETM is the first framework that effectively integrates GFMs for scATAC-seq analysis and can generalize to both unseen cells and peaks via the VAE-encoder and GFM architecture design. This led to competitive performance on downstream transfer learning tasks in a wide range of scATAC-seq datasets, including cross-tissue, cross-species, and cross-modality transfer learning. Our work GFETM presents a novel solution to advance the representational learning of cells based on their epigenomic profiles and to deepen our insights into the genomic elements that are associated with cell-type-specific chromatin accessibility.

A notable capability of GFTEM is its proficiency in transfer learning. With single-cell ATAC-seq datasets often limited in scope across many studies, the utilization of extensive, relevant reference datasets becomes critical. GFETM’s effectiveness in learning sequence information enables superior performance in transfer learning across diverse tissues, species, and sequencing modalities. This feature addresses the challenge of limited data availability, allowing for the application of models trained on reference datasets to new datasets containing unseen cells and peaks. Moreover, the interpretability of GFETM makes it stand out among the variety of black-box deep-learning-based tools developped for genomics research. In contrast to methods that produce outputs with abstract statistical interpretations, GFETM generates topics with direct and intuitive biological relevance. These topics frequently align with transcription factor (TF) motifs, providing insights into gene regulation. This interpretability offers a deeper understanding of the epigenomic landscape’s role in influencing cellular states and identity.

For future work, we will explore more efficient fine-tuning strategies on the GFM and other transfer learning and domain adaptation techniques [45, 46]. We will investigate the impact of single nucleotide polymorphism (SNPs) on the cell-specific chromatin accessibility via in-silico mutageneiss or more sophisticated explainable AI methods [47]. We will apply GFETM to disease-specific scATAC-seq data including Myocardial Infarction [48] and Alzheimer’s Disease [49] and explain the missing heritability of the corresponding genome-wide association studies. Lastly, extending GFETM to multi-omic modeling of scATAC-seq with scRNA-seq is also a promising venue [50].

In summary, we apply the principals of LLMs to the text-like structure of DNA, facilitating a nuanced exploration of the chromatin landscape at the single-cell level. Such a framework promises to enhance the research community’s ability to decode complex cellular processes, by interpreting chromatin remodeling with the same acuity that LLMs process semantic information. The implications of GFETM in epigenomic research is promising as it enables scientists to navigate the vast genomic datasets with foundation models pre-trained on millions of genomic sequences, which can shed novel insights into the gene regulation that dictate diverse cell functions and dysfunctions.

## 4. Methods

### The ETM component

We model the scATAC-seqs data as a cells-by-peaks matrix **X** ∈ {0, 1}*^N×M^* with *N* cells and *M* peaks. We model the peak data using multinomial distribution with the expected peak rate parameterized as **R** ∈ [0, 1]*^N×M^*. We decompose **R** into cells-by-topics θ ∈ [0, 1]*^N×K^* and topics-by-peaks ***β*** ∈ [0, 1]*^K×M^* (i.e., **R** = θ***β***). The two matrices are softmax-normalized over *K* topics (i.e., columns) and *M* peaks (i.e., rows), respectively. Specifically, for each cell *c* ∈ {1*, …, N* }, the cell topic mixture is ***θ****_c,._* such that 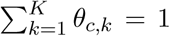; for each topic ***β****_k,._*, where *k* ∈ {1*, …, K*}, 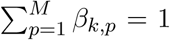. We further decompose the topic distribution into the topic embedding ***α*** ∈ ℝ*^K×L^* and peak embedding ***ρ*** ∈ ℝ*^L×M^*, where *L* denotes the size of the embedding space. Thus, the probability of a peak belonging to a topic is proportional to the dot product between the topic embedding and the peak embedding (***β****_k,._* ∝ ***α****_k,._****ρ***). Formally, the data generative process of each scATAC-seq profile of cell *c* ∈ {1*, …, N* } can be described as follows:

1. Draw a latent cell type mixture ***θ****_c_* for a cell *c* from logistic normal ***θ****_c_* ∼ LN (0, **I**):

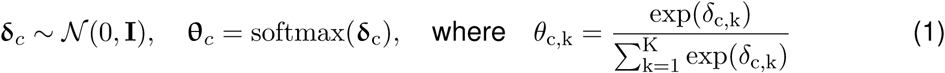
2. For each peak token *w_c,i_*, where *i* ∈ {1*, …, N_c_*} among the *N_c_* peaks observed in cell *c*, draw a peak index *p* ∈ {1*, …, M* } from a categorical distribution Cat(**r**_c_):

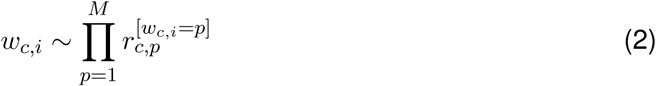

where 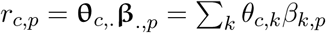 denotes the categorical rate for cell *c* and peak *p* and

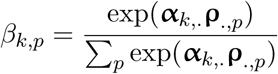

Given **x***_c_*∈ [0*, N_c_*]^1*×M*^, as the vector of the peak count over all *M* peaks, the likelihood over the *N_c_* peak tokens *w_c,i_*’s follows a multinomial likelihood:

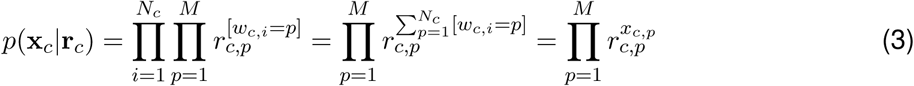

When integrating multiple scATAC-seq datasets, in order to account for batch effect, the chromatin accessibility rate *r_c,p_* is further parameterized as:

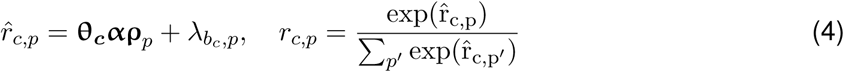

where 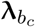 is an optional parameter which depends on the batch index *b_c_* ∈ {1*, …, B*} of cell *c* for B batches.

The log marginal likelihood is approximated by an evidence lower bound (ELBO):

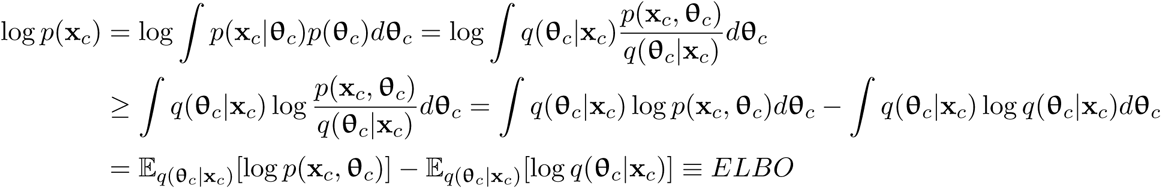

where *q*(***θ****_c_*|**x***_c_*) is a proposed distribution to approximate the true posterior distribution *p*(***θ****_c_*|**x***_c_*). Maximizing the ELBO minimizes the Kullback-Leibler (KL) divergence between the proposed and true posterior distributions: 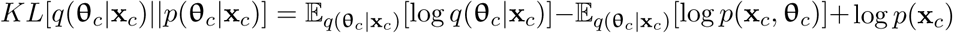 because they sum to the constant log marginal likelihood (i.e., *ELBO*+*KL*[*q*(***θ****_c_*|**x***_c_*)||*p*(***θ****_c_*|**x***_c_*)] = log *p*(**x***_c_*)).

Using the variational autoencoder (VAE) framework [51], the variational distribution is defined as *q*(***δ****_c_*|**x***_c_*) = N (***δ****_c_*; ***µ****_c_,* diag(***σ***_c_)), where ***µ****_c_* and ***σ****_c_* are the outputs of a neural network function (i.e., the encoder): [***µ****_c_*, log ***σ****_c_*] = *f* (**x̂***_c_*|**W**). Here **x̂***_c_* is the normalized Term Frequency – Inverse Document Frequency (TF-IDF) count [52]. Finally, the ELBO is further divided into the marginal log multinomial likelihood under the variational distribution and the regularization term, which is the KL divergence between the variational and the prior distributions: 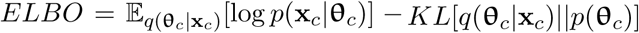. Given the standard Normal prior, the regularization term has a closed-form of 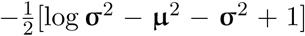. The variational likelihood is approximated by Monte Carlo, where we use sampled ***δ̃***_*c*_ ∼ ***µ****_c_* + ***σ****_c_*N (0, **I**) from the proposed distribution to evaluate the likelihood: 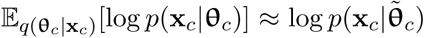. Note that ***θ̃***_*c*_ is deterministic given ***δ̃***_*c*_ (Eq. 1). Training of the encoder network is done by stochastic gradient descent.

### The GFM component

We use a pre-trained GFM to extract peak nucleotide sequence embeddings. Specifically, given input peak *p* containing the chromosome index and start/end position information, we extract the corresponding DNA sequence from the reference genome. The sequence is then tokenized into input tokens by specific tokenization algorithms in specific GFM. The input tokens are then fed into the pre-trained GFM to obtain the token embedding. The sequence embedding ***ρ****_p_* for each peak is computed by average pooling the token embeddings. In this study, we use existing released GFMs including DNABERT [21], Nucleotide Transformers [23], DNABERT-2 [22] and HyenaDNA models [24]. We include different versions (i.e. kmer, length, size) of these GFMs for comprehensive evaluation. Specifically, DNABERT [21] is based on the BERT [53] architecture and the DNA sequences are tokenized using k-mer tokenization. DNABERT-2 [22] is also based on the BERT but uses more advanced techniques including Flash-Attention [54] and Attention with Linear Biases [55] to further improve the training efficiency and model performance. Byte Pair Encoding [56] is also used as the tokenization algorithm for DNA sequences tokenization. Nucleotide Transformers are also based on the BERT architecture and k-mer tokenization, but trained on not only human reference genome but also 3000 individual genomes from 1000 Genomes Projects as well as 850 species genomes with the largest parameter size up to 2.5B. HyenaDNA is a fast and lightweight GFM based on long implicit convolutions with Fast Fourier Transform for sub-quadratic time complexity to scale to ultralong sequences at the single-nucleotide resolution. Detailed information on the included models is shown in Supplementary Table S1.

### Leveraging the peak embedding from GFM in ETM

We extracted the sequence embeddings from the last layer of the pre-trained GFM for all the peaks in the dataset and concatenate the embeddings into the peak embeddings ***ρ*** ∈ ℝ*^L×M^* in the ETM component. We experimented three strategies of leveraging these peak embeddings of GFM. In the first strategy, the peak embeddings from the last layer of GFMs were used as they are. In the second strategy, we use the peak embedding to initialize the model parameters ***ρ*** in the ETM component and jointly update it with the other ETM parameters during the training. We experimented with different GFMs and reported the performance from fixing and fine-tuning the peak embeddings in Fig. 2.

However, directly using the peak embedding as the model parameters result in too many learnable weights for large number of peaks, which is the case in scATAC-seq atlas. To further improve the model performance, we developed a joint-training and fine-tuning strategy for the ETM and GFM components, respectively. Specifically, we fine-tuned the last two transformer layers from GFM together with the ETM component. This allows the GFM and ETM to jointly optimize the ELBO objective function. Specifically, we modified Eq 4 to amortize the peak embedding learning on the GFM:

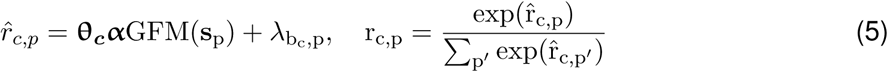

where GFM(**s**_p_) is the output of the last layer of the GFM for the peak embedding ***ρ****_p_* based on the one-hot sequence matrix **s***_p_*. This requires more computational cost compared to the first way. Therefore, we adopted a minibatch peak sampling strategy. Given a minibatch of cells, we first randomly sample *M ^′^* ≤ *M* peaks among these cells and obtain the nucleotide sequence embedding 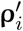 of those *M ^′^* peaks through forward pass of the GFM. The sequence embedding 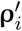 from the GFM are then multiplied with the latent topic mixture ***θ*** and topic embedding ***α*** from the ETM (Eq 4) to obtain the expected transcription rate.

In practice, to reduce the computational costs of joint-training, we sample 384 peaks in each iteration (*M ^′^*=384). As some of GFMs including DNABERT-2 and Nucleotide Transformers contain very large parameters, integrating them with ETM to perform joint-training is hard considering the resource constraints. Therefore, we opted pre-trained DNABERT because it is fairly light weighted and based on standard transformer architecture.

### Transfer learning of GFETM

We performed transfer learning of GFETM in zero-shot and continual training settings. In the zero-shot transfer setting, the GFETM model was first trained on a source scATAC-seq dataset. The model parameters were then frozen and the model was tested on the target scATAC-seq dataset from different species or tissues. In the continual training setting, the GFETM model was first pretrained on the source scATAC-seq dataset and then fine-tuned on the target scATAC-seq dataset from different species or tissues. The model was evaluated on the target scATAC-seq dataset.

Note that ATAC-seq peaks in different species or experiments are different. Therefore, we aligned different peaks from different datasets to enable the transfer learning. Concretely, We utilized LiftOver (http://genome.ucsc.edu/cgi-bin/hgLiftOver) for the conversion of genome coordinates from the mouse reference genome mm9 to the human reference genome hg19, facilitating the alignment of peaks between mouse and human to enable cross-species transfer learning on scATAC-seq data. We omitted those peaks that failed the conversion. When aligning peaks between datasets, we considered peaks *i* from dataset 1 and peak *j* from dataset 2 aligned if there is overlap between them. The overlapped peaks were then used to represent the features for the two aligned datasets. Peaks that do not have overlap between the datasets were ignored for the transfer learning experiments.

In the cross-omic transfer setting, we collected the multiome data with scRNA and scATAC seq measured in the same cells from Chan Zuckerberg CELLxGENE. The scRNA and scATAC-seq datasets were aligned on a common gene basis by the dataset curator [36, 48, 57] (through Cell-Ranger https://github.com/10XGenomics/cellranger). We extracted the DNA sequences −/+ 256 nucleotide of the TSS of each gene as the peak sequences based on the GENCODE annotation version 43 (gencode.v43.basic.annotation.gtf). In the cross-omic transfer learning, based on the aligned features, we first trained the model on the scRNA-seq and scATAC-seq dataset and then transferred to modeling the scATAC-seq and scRNA-seq dataset, respectively.

### Generalization to unobserved peaks

Given the GFETM trained on an scATAC-seq dataset using *N* cells and *M* peaks from all but one chromosome, we assess the accuracy of inferring the *M ^′^* unseen peak accessibility on the held out chromosome for each individual cell.

Specifically, imputing unseen peaks takes 3 stages (Supplementary Fig. S11). In stage a, we trained the original GFETM on the *N* cells and *M* peaks, where the reconstruction loss corresponds to the multinomial likelihood: 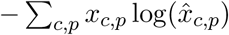, where the reconstructed peak is normalized with softmax across peaks 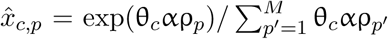. The goal of this stage is to learn a good encoder and a reasonably good decoder for the particular data at hand. In stage b, we freeze the GFETM encoder and continue to train the GFETM decoder including the topic embedding α and the GFM with negative Bernoulli log likelihood (i.e., cross-entropy loss) as the new reconstruction loss for the *M* peaks: 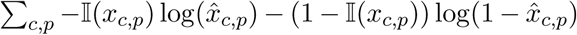, where Ι(*x*) is an Boolean indicator function that returns 1 with *x >* 0 and 0 otherwise, and *x̂_c,p_* = *σ*(θ*_c_*αρ*_p_*) with *σ*(*x*) being the sigmoid function. For this phase, we switch to the cross entropy loss to improve the specificity of the imputation especially for sparse signal at each individual cell. In stage c, (i.e., the inference stage), for each cell *c* ∈ {1*, …, N* } and *M* peaks seen in the training data, we first use the trained GFETM encoder to generate the cell embeddings θ*_c_*. To impute whether the cell contains the *M ^′^* new peaks from the held-out chromosome, we feed the corresponding *M ^′^* DNA sequences to the trained GFM from stage b, which generates the peak embeddings ρ*^′^* ∈ *R^L×M′^*. Finally, we compute the scores for the *M ^′^* peaks as the dot product of the cell embedding, topic embedding, and peak embedding: θ_*c*_αρ*^′^* ∈ *R*^1^*^×M′^*.

### Marker gene enrichment analysis on denoised matrix

The *R* = {*r_c,p_*} matrix with 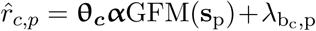 and 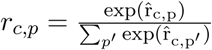 can be considered as a denoised version of the original scATAC-seq matrix. We evaluated whether the denoised matrix has higher enrichment in identifying cell type specific marker genes. Given the raw data **X**, the denoised matrix *R* by GFETM or scBasset, we performed Wilcoxon test to detect marker peaks for each cell type (i.e., comparing peak signals among cells in one cell type against the cells in other cell types). We selected marker peaks at Bonferroni adjusted p-values or false discovery rate (FDR) *<* 0.1 to correct multiple testing over all peaks and log-fold changes larger than 0. To assess whether the candidate marker peaks are biologically meaningful, we annotated them by the in-cis genes if the peaks are within a defined genomic distance upstream from the TSS of the corresponding genes. We experimented different FDR threshold, log-fold changes threshold, and the TSS distance in terms of the enrichment for the known marker genes of the cell type from CellMarker 2.0 database http://bio-bigdata.hrbmu.edu.cn/CellMarker/. We computed the overlap between cell type marker genes and the in-cis genes downstream of the marker peaks identified by the raw data and the denoised matrix. To assess the statistical significance of the overlap, we performed hypergeometric test.

### Motif Scoring Analysis

Estimating the motif activity from the scATAC-seq datasets is an important function of GFETM. Concretely, we computed the TF motif embeddings GFM(**m**) by feeding the motif sequences to the GFM component of trained GFETM. The dimension of cell embeddings ***θ****_c_* from GFETM is *K* (topic size) while the dimension of motif embeddings is *L* (embedding size). Usually *K* is much smaller than *L*. Therefore, to estimate the TF motif activity score, we multiplied the cell embeddings ***θ****_c_* by the topic embeddings ***α*** to match the dimension of motif embeddings. The activity of cell *i* on TF *j* is computed as the cosine similarity between (***θ****_c_****α***)*_i_* and GFM(**m***_j_*). We used a human cortex multiomics profiling dataset [32] containing 8981 cells with paired scRNA-seq and scATAC-seq profiles. We used the TF motifs collected by [32] which are validated to be relevant to the transcription regulation of human cortex: PAX6, INSM1, SOX9, TCF4, TCF3, TFAP2C, FOS, NEUROD2, BHLHE22, JUND, MEF2C, POU2F2, NFIA, MEIS2, EOMES. The correctness of the motif scoring output is evaluated by the correlation between the estimated motif activity with the gene expression of the corresponding TF gene following [18]. We compared GFETM with CellSpace and scBasset. It is worth noticing that scBasset, which encodes the peak DNA sequences with a CNN, takes multiple background sequences with or without the target TF motif sequence to estimate the motif activity, while GFETM can directly encode the short TF motif sequences, which is much more efficient.

### Attention Analysis of GFM

The attention mechanism of GFM provides valuable insights on the latent interactions between different regions inside the GFM. Hence, we analyzed the attention mechanism of GFM to investigate whether GFM learns the intricate activities of transcription factors within the peak regions of scATAC-seq datasets. Concretely, given a peak sequence, we extracted the last layer attention of the GFM component from GFETM and used max pooling to incorporate information from different attention heads. We matched the TF motif regions on the peak sequences with the function Bio.motif.pssm.search from the biopython package with matching score threshold set to 5 ^1^. We aggregated the attention weight based on the TF motif regions.

### Differential analysis of topics mixtures

For a given topic *k*, we computed the difference of average topic scores between each cell type or condition and the rest of the cell types or conditions. To assess the statistical significance, we conducted *N* = 100, 000 permutation tests by randomly shuffling the label assignments among cells and recalculating the average topic score difference based on the resulting permutations. The empirical p-value was computed as (*N ^′^* + 1)*/*(*N* + 1), where *N ^′^* represents the number of permutation trials where the difference exceeded the observed value. We then applied Bonferroni correction by multiplying the *p*-value by the product of the topic number and the number of labels. Topics that are below Bonferroni-corrected q-value = 0.1 were deemed DE topics with repect to the cell type or condition.

### Motif extraction and enrichment analysis

Given the trained ETM model and the topic-by-peak matrix (i.e., ***β*** ∝ ***αρ***), we extracted the top 100 peaks for each topic. Using the sequences of these top peaks as query, we performed motif enrichment analysis to identify the known motifs that are enriched in the peak sequences using the Simple Enrichment Analysis (SEA) pipeline [58] from the MEME suite [59]. SEA utilizes the STREME motif discovery algorithm [60] to identify known motifs that are enriched in input sequences. We used the HOmo sapiens COmprehensive MOdel COllection (HOCOMOCO) Human (v11) and HOCOMOCO Mouse (v11) motif database [61]. Motifs with E-values ≤ 10 were identified as enriched motifs.

### Correlating cell-type similarity and transfer learning performance

We computed the Jaccard similarity between two tissues *A* and *B* in terms of the number of shared cell types. For each target tissue, we then ranked all the source tissues based on their similarity with it. In parallel, we ranked all the source tissues by its zero-shot transfer performance to the target tissue in terms of the ARI on the target tissue. We used Kendall’s tau *τ* and Spearman rank-order correlation coefficient *ρ* to measure the correspondence between two rankings. The cell type distribution of the dataset is shown in Supplementary Fig. S17 and S19

### Data preprocessing

We used the Human HSC Differentiation hematopoietic dataset [62], 10x multiome PBMC dataset, and 10x mouse brain dataset to benchmark the model performance following [7]. The Human HSC Differentiation dataset contains scATAC-seq data of human hematopoiesis of 10 hematopoietic cell types, with 2034 cells and 6 batches included in total. The 10x multiome PBMC dataset contains cryopreserved human peripheral blood mononuclear cells (PBMCs) of a healthy female donor aged 25 obtained by 10x Genomics, with 2714 cells included in total. The 10x mouse brain dataset contains cells obtained from fresh embryonic E18 mouse brain, with 4878 cells in total. To evaluate atlas-level integration and transfer learning, we used the Catlas-human dataset [63], which contains 1.3 million cells, and we used the Cusanovich-mouse dataset [12], which is a large scale multi-tissue mouse scATAC-seq data. To discover disease-specific topics and perform cross-omic transfer, we use the human kidney diabetic dataset [36], human myocardial dataset [48], and human ovary dataset [57].

For Human HSC Differentiation dataset, 10x multiome PBMC dataset and 10x multiome mouse brain dataset, we followed the same data preprocessing steps in [20]. Peaks accessible in fewer than 1% cells were filtered out. For cross-species transfer learning, we selected heart, lung and intestine from Cusanovich-mouse and Catlas-human dataset and sampled 5000 cells from each tissue. To perform cross-tissue transfer learning, we selected 12 mouse tissues from Cusanovich-mouse and 21 human tissues from Catlas-human and sampled 2000 cells from each tissue. This was to maintain the same size per tissue and to include as many tissues as possible because some tissues contain fewer than 5000 cells. We kept 2^16^ highly variable peaks in the dataset. For RNA-ATAC transfer learning, we sampled 10,000 cells from the scRNA-seq data and scATAC-seq data respectively.

### Details in GFETM Training

The encoder of the GFETM is an 2-layer MLP with hidden size 128 with ReLU activation and batch normalization and outputs the ***µ*** and ***σ***. The dimension of peak embedding ***ρ*** was set to match output embedding size of the pretrained GFM (i.e. 768 for DNABERT). We optimized the model with Adam optimizer with learning rate 0.005 for both ETM and GFM components. We also added *β* hyperparameter to weight the regularization (i.e., *β*-VAE [64]). As a high weight penalty of *β* on the KL divergence tends to produce poor cell embedding, we used a small weight penalty *β* = 1*e* − 6 on the KL divergence. The impact of several hyperparameters on the model performance is shown in Supplementary Material S1.4.

### Baseline Methods

We followed the evaluation settings as described in [7,20]. We used the results for several baseline methods in [20]. For SIMBA [19], we used the default settings. For CellSpace [18]. We trained the Cusanovich-mouse dataset using the default model parameters as suggested in the CellSpace paper and documentation. The 10x mouse brain dataset was trained with the sampled sequence length parameter set to 300 base pairs as was done in the CellSpace paper for a similarly sized dataset. We performed extensive tuning and selected the training epoch that generated the highest performance (epoch 40) for the mouse brain dataset.

## 5. Data availability

No new single-cell ATAC-seq datasets are generated in this study. All the datasets used in this study are public available. We collected Human HSC Differentiation dataset (GSE96772) [62] PBMC dataset (https://support.10xgenomics.com/single-cell-multiome-atac-gex/datasets/2.0.0/pbmc_granulocyte_sorted_3k), Mouse brain dataset (https://support.10xgenomics.com/\single-cell-multiome-atac-gex/datasets/2.0.0/e18_mouse_brain_fresh_5k), Catlashuman (GSE184462) [63], Cusanovich-mouse (GSE111586) [12], Myocardial Dataset [48], Ovary Dataset (EGAS00001006780) [65] and Kidney Diabetics (GSE195460) [36].

## 6. Code availability

Source code of this study has been deposited in https://github.com/fym0503/GFETM.

## 7. Competing interest statement

The authors declare no competing interests in this study.

## 8. Acknowledgments

Y.L. is supported by Canada Research Chair (Tier 2) in Machine Learning for Genomics and Healthcare (CRC-2021-00547) and Natural Sciences and Engineering Research Council (NSERC) Discovery Grant (RGPIN-2016-05174). A.O. is supported by a Fonds de Recherche du Québec Santé (FRQS) training scholarship. This work was partially supported by CIHR PJT-180505, FRQS 295298, 295299, and NSERC RGPIN-2022-04399 to JD.

## 9 Author contributions

Y.L. and J.D. conceived the study. Y.L., J.D., and Y.F. developed the methodology. Y.F. implemented the software and conducted the experiments under supervision of Y.L. and J.D.. A.O ran several baseline methods. Y.F., Y.L., and J.D. analyzed the results. Y.F., Y.L., and J.D. wrote the initial draft of the manuscript. All of the authors reviewed and wrote the final version of the manuscript.

## S1 Supplementary Material

### S1.1 Comparison with CNN-based ETM

To demonstrate the superiority of GFM over the traditional CNN architecture, we built a baseline that replaces GFM with CNN. The architecture of the CNN baseline is shown in Supplementary Fig. S2. We benchmarked the performance of this ETM+CNN baseline and the proposed GFETM method on several settings in Supplementary Fig. S3. We observed that GFETM outperforms ETM+CNN on common benchmarking datasets (Buen18 and Mouse brain datasets) and crosstissue transfer-learning setting (Cerebellum to Brain and Lung to Heart) in terms of ARI. It is important to note that there are many ways of designing the CNN architectures that may work better on this task, which we have not thoroughly explored to reach this conclusion.

### S1.2 Scalability in terms of running time and memory

We recorded the runtime per epoch as an average over 100 epochs. GFETM increases slowly with the increase of peak numbers and scales linearly with the number of cells (Supplementary Fig. S9c). As scBasset takes peak sequences as input and the cell embeddings are trainable parameters in the model, the runtime of scBasset scales linearly with peak number and only slightly increases with the number of cells.

Regarding the total runtime, on the dataset with 20k cells and 20k peaks, scBasset took around 5500 seconds to converge (i.e., 1.5 hrs) and GFETM took around 15000 seconds (i.e., 4 hrs). Both are acceptable considering the dataset scale. The total runtime of GFETM also scales linearly with the number of cells in the dataset as the input mini-batch is based on cells while the total runtime of scBasset scales linearly with the number of peaks as the input mini-batch is based on peaks. As GFETM is based on GFMs, which has much larger parameters than CNN-based scBasset (88 million parameters for GFETM and 5 million parameters for scBasset), the runtime of GFETM is expected to be higher. If using several GPUs for computation, the computational efficiency of GFETM may be further improved.

### S1.3 Reconstruction on training and testing peaks

As reconstruction of the scATAC-seq peak accessibility data is the primary training objective, we validated that GFETM can reconstruct the scATAC-seq peak accessibility accurately and generalize well on new cells and peaks. As shown in Supplementary Fig. **??**a, first, we observe no correlation at the initialization stage; after training for 100 epochs, the reconstruction and observed were weakly correlated; after training for 1000 and 2000 epochs, the correlation became strong, which demonstrates the effectiveness of the training procedure.

We then evaluated the generalization of chromatin accessibility prediction across the training and testing cells. As shown in Fig. **??**b top panel, for the test cells, the predicted accessibility also exhibit high correlation with the groundtruth. Thus, our method is generalizable in terms of chromatin accessibility prediction on training and testing cells.

We then evaluated the GFETM’s ability to model unseen peaks in terms of reconstruction. First, we trained the GFETM model on a dataset with *N* cells and *M* peaks. After the training, the trained ETM parameters were discarded and only the GFM parameters were kept. We then trained a new GFETM model with its GFM parameters inherited from the previously trained GFM on the dataset with *N* cells and *M* + *M^′^* peaks. To assess the generalizability of the GFM, we froze its parameters during the training on the new *M* + *M^′^*-peak dataset. Indeed, as shown in Supplementary Fig. **??**c, in the beginning of the training, the performance on seen peaks is better than unseen peaks. With the progress of training, the performance on imputing the unobserved peaks becomes gradually better and eventually matches the performance on reconstructing the observed peaks, which demonstrates the generalizability of the GFM.

### S1.4 Hyperparameter tuning of GFETM

GFETM is generally robust to different hyperparameter setting and configurations. We mainly experimented with two important hyperparameters, the version of DNABERT and the number of topics. First, there are four versions of DNABERT released by the authors of DNABERT, including 3-mer, 4-mer, 5-mer and 6-mer. We used the 6-mer version of the DNABERT as this is the version used throughout the original paper and github code release. As shown in Fig. S4, we ran the experiments 5 times with different random seeds. We observe that different versions of DNABERT and different topic numbers do not have significant impact on the performance of our model, which demonstrates the robustness of our method.

### S1.5 Supplementary Figures

**Supplementary Figure S1:**
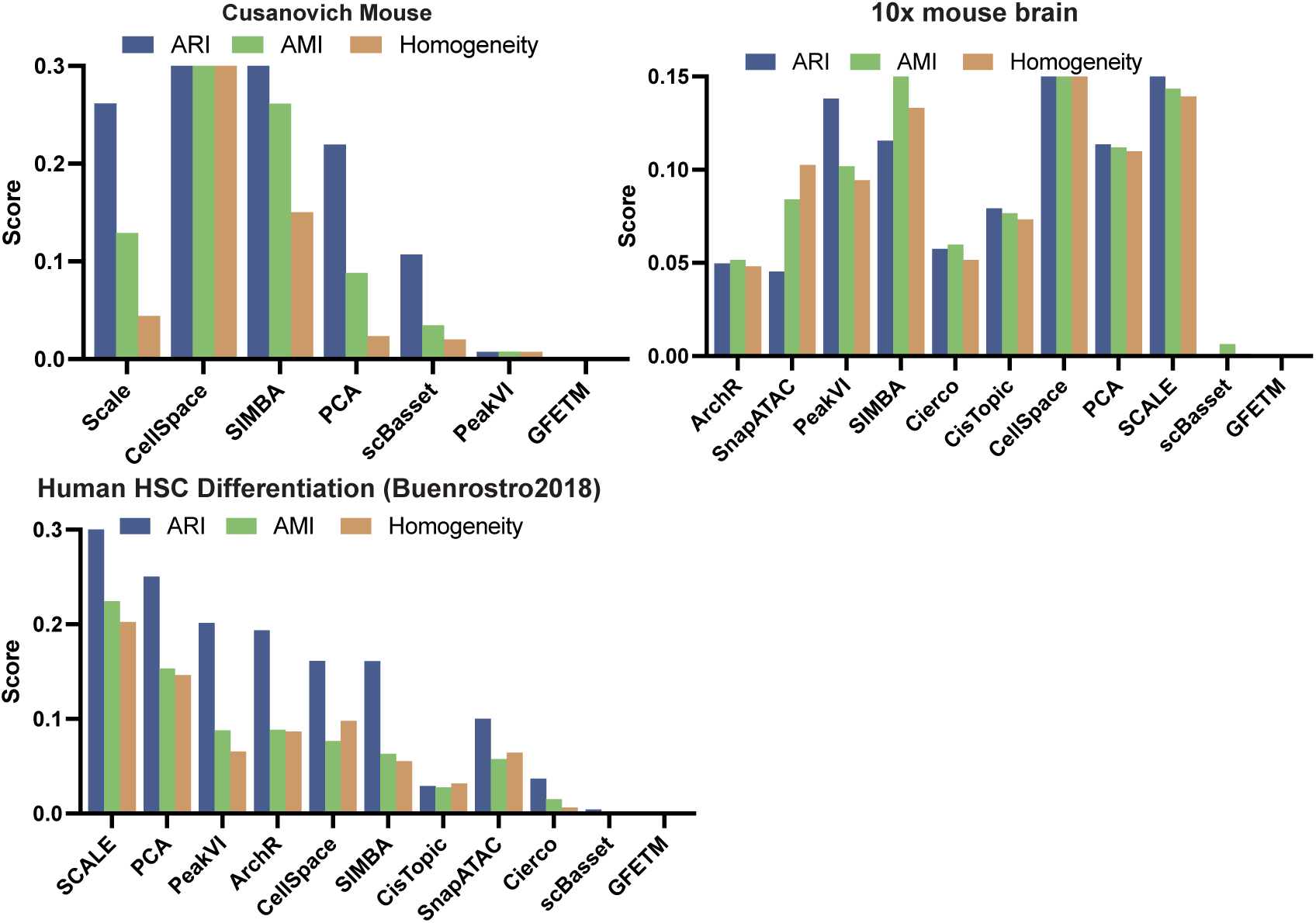
GFETM’s improvements over the existing methods. We experimented each method on three datasets namely Human HSC Differentiation, 10x mouse brain and Cusanovich Mouse in terms of ARI, AMI and Homogeneity. The bars for each method are calculated the its score subtracted from the score achieved by the GFETM. Therefore, a positive score indicates an improvement of GFETM over that method.

**Supplementary Figure S2:**
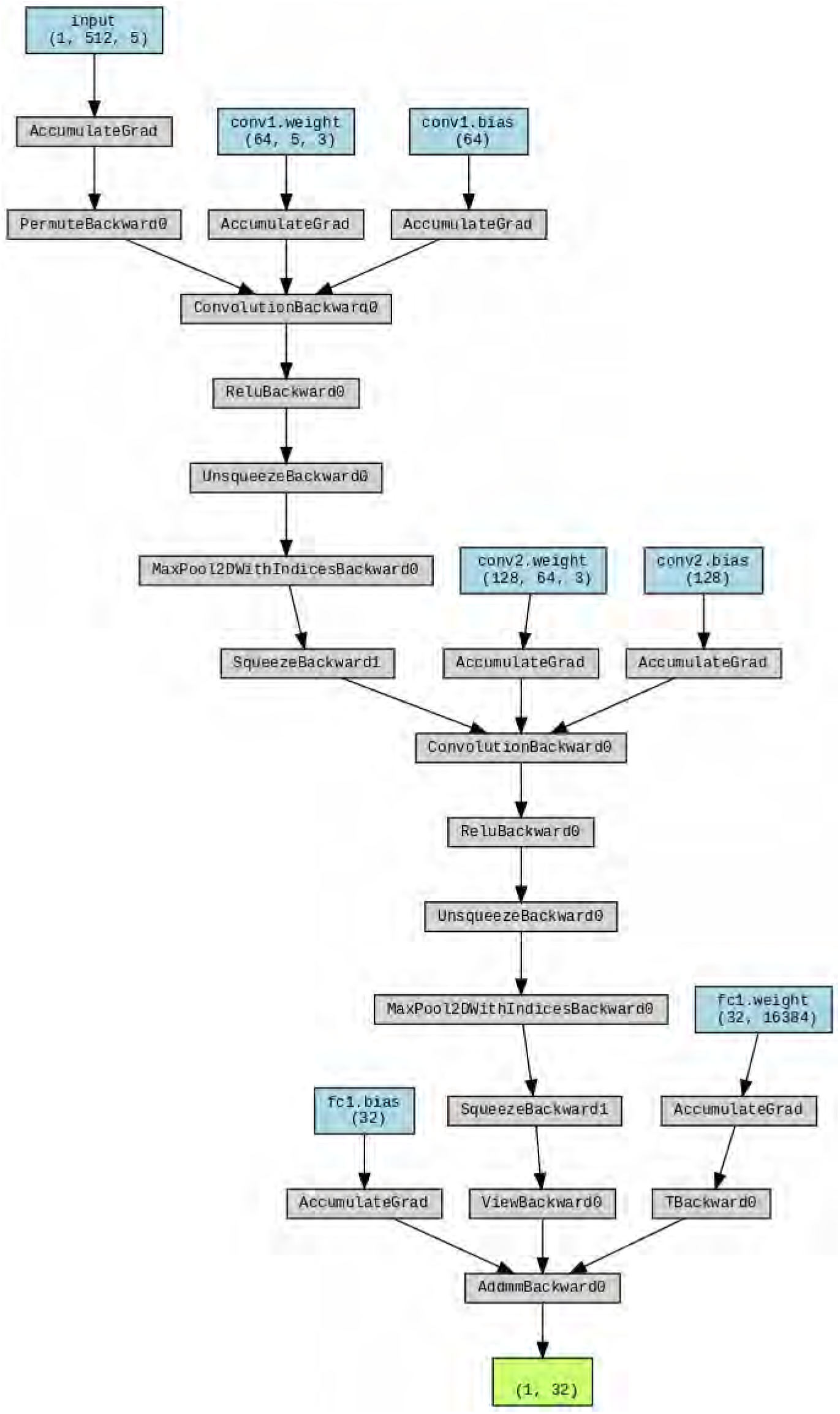
Architecture of the CNN baseline that is used to replace the DNA sequence Transformer. The model architecture is generated using torchviz.

**Supplementary Figure S3:**
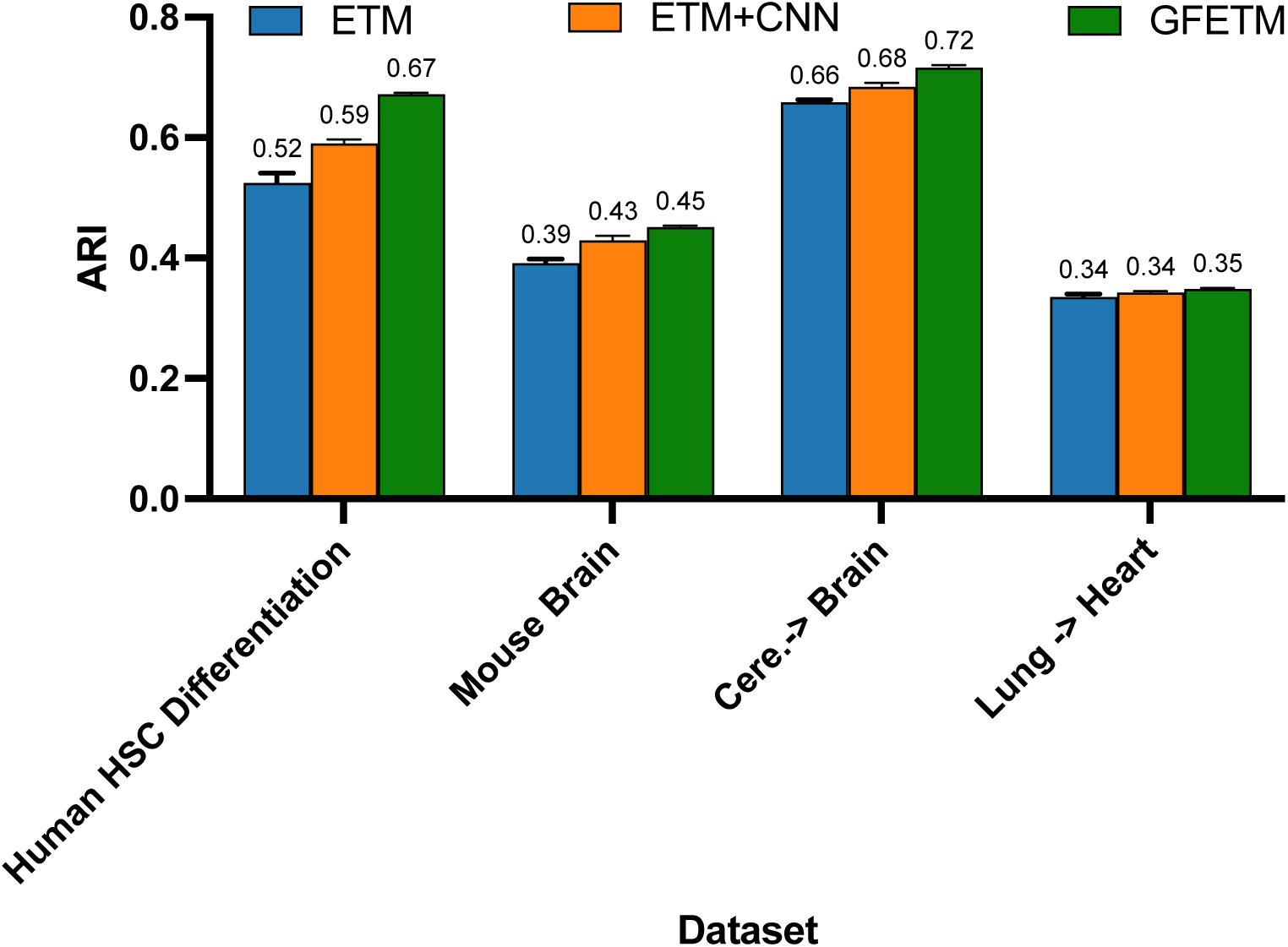
Performance Comparison between the ETM, ETM+CNN baseline and the proposed GFETM on several settings. Cere. stands for Cerebellum

**Supplementary Figure S4:**
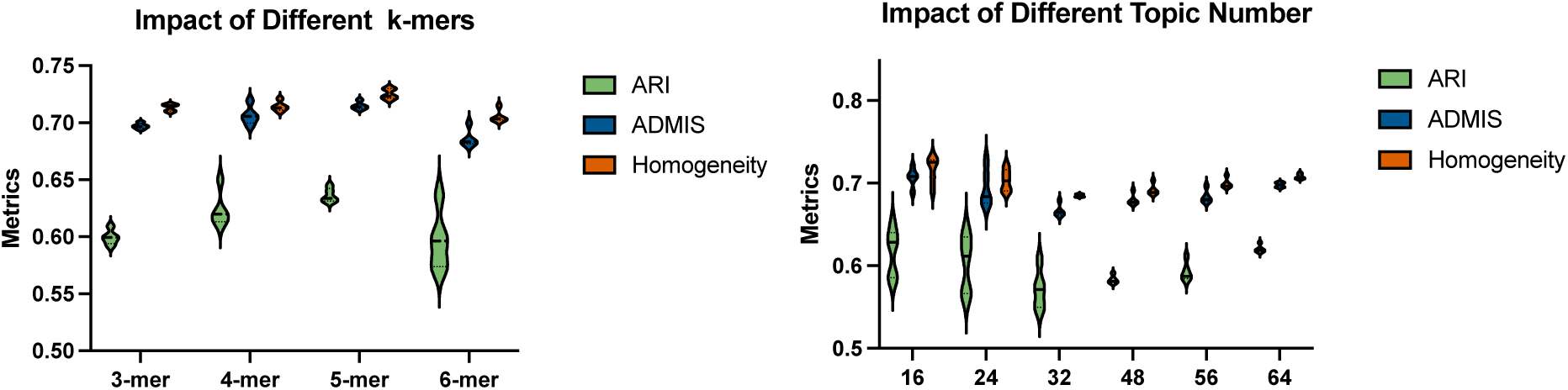
Left: impact of k-mers from different versions of DNABERT model on the model performance. Right: impact of different numbers of topics on the model performance.

**Supplementary Figure S5:**
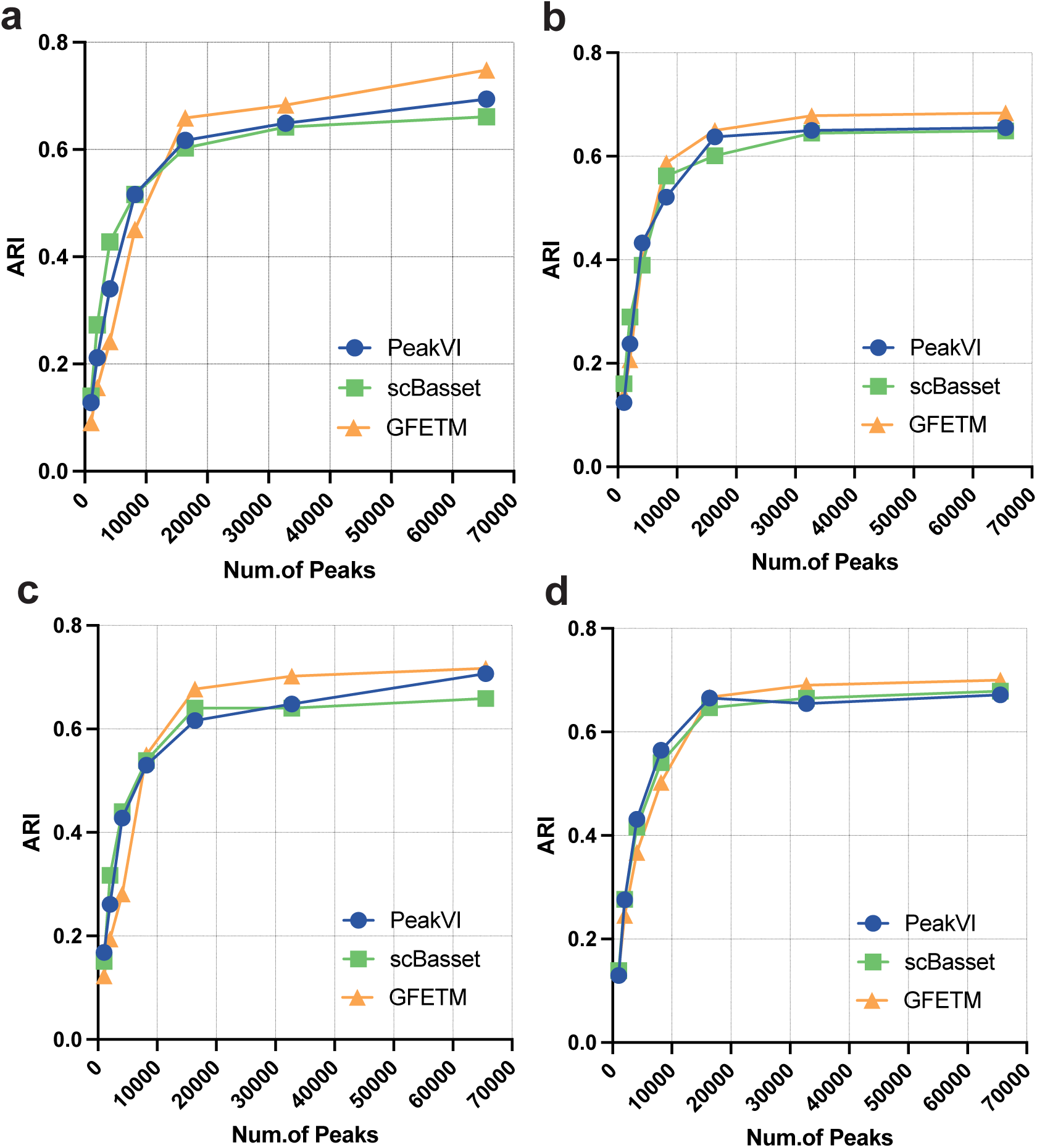
Scalability on large-scale datasets. scBasset and PeakVI are used as baseline methods. a. the panels indicate the results for 20000 cells from the Cusanovich-Mouse dataset b. 40000 cells c.60000 cells d. 80000 cells.

**Supplementary Figure S6:**
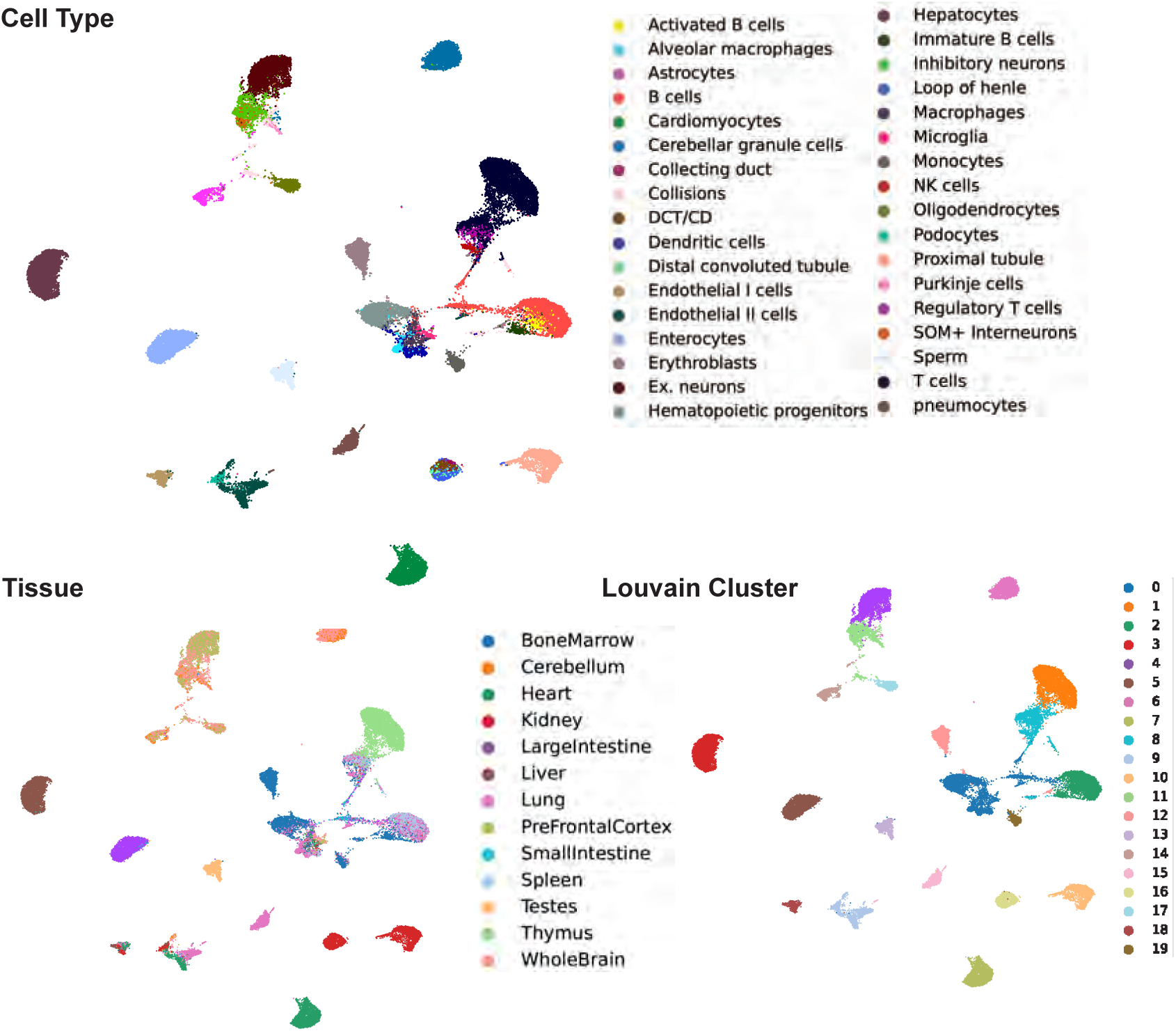
Visualization of GFETM latent embeddings on Cusanovich-mouse atlas-level datasets across cell types, tissues and predicted labels.

**Supplementary Figure S7:**
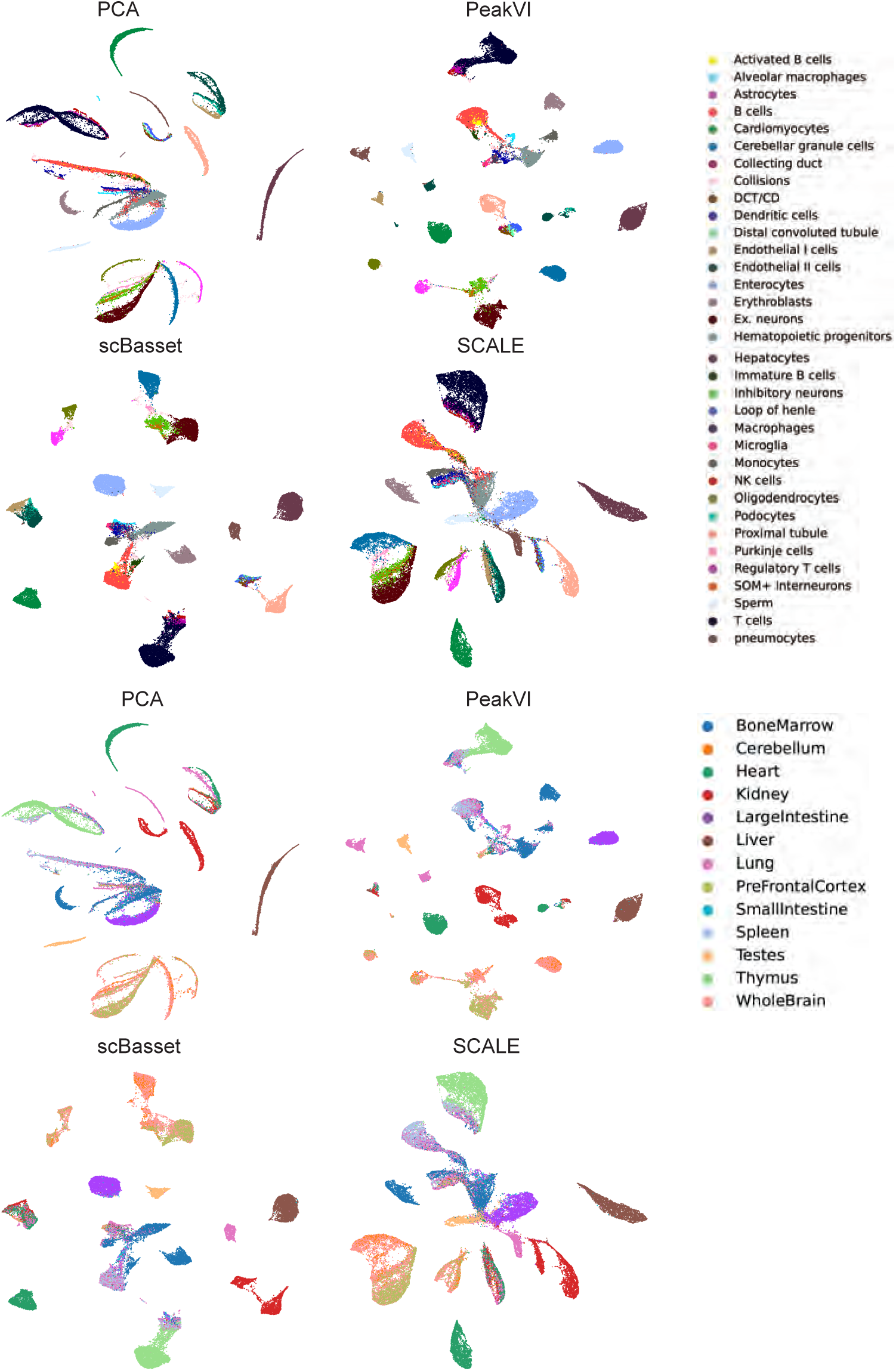
Visualization of latent embeddings on Cusanovich-mouse atlas-level datasets across cell types, tissues of baseline methods.

**Supplementary Figure S8:**
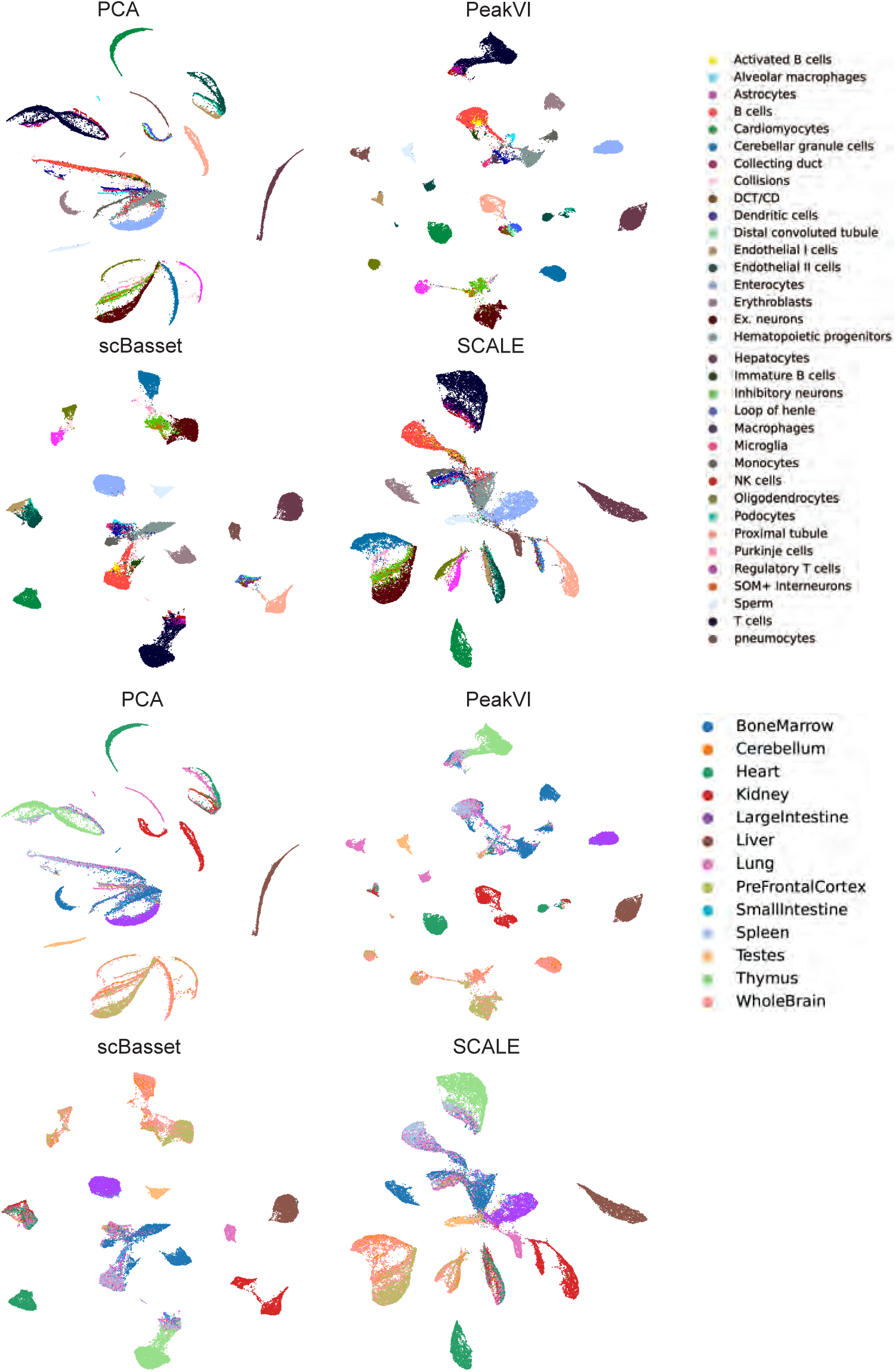
Visualization of latent embeddings on Cusanovich-mouse atlas-level datasets across predicted labels of baseline methods.

**Supplementary Figure S9:**
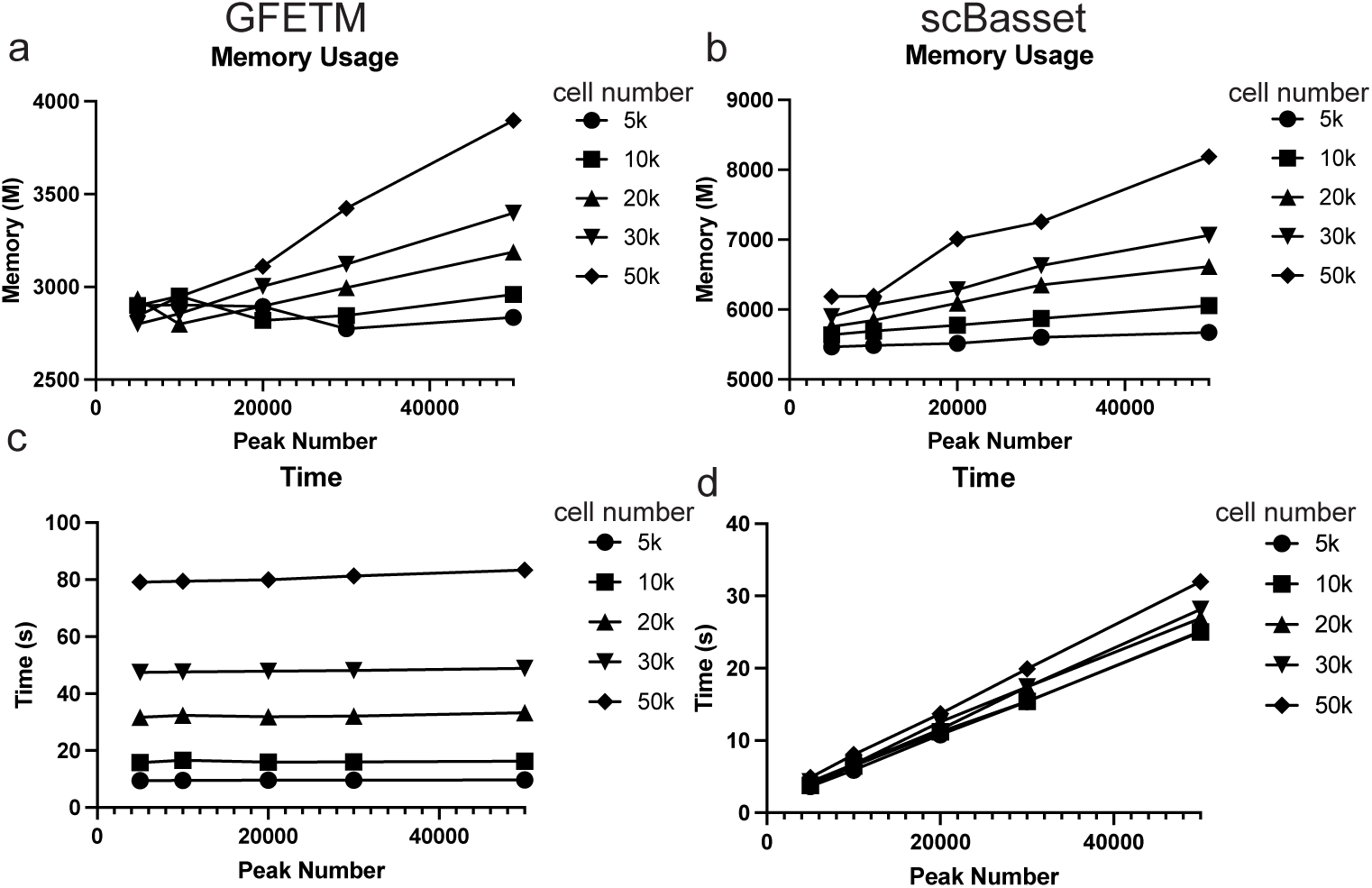
Memory and runtime analysis of GFETM and scBasset. a & b. Memory usage of GFETM and scBasset with respect to cell number (scatter point shapes) and peak number (x-axis). c & d. Runtime (per epoch) of GFETM and scBasset with respect to cell number (scatter point shapes) and peak number (x-axis).

**Supplementary Figure S10:**
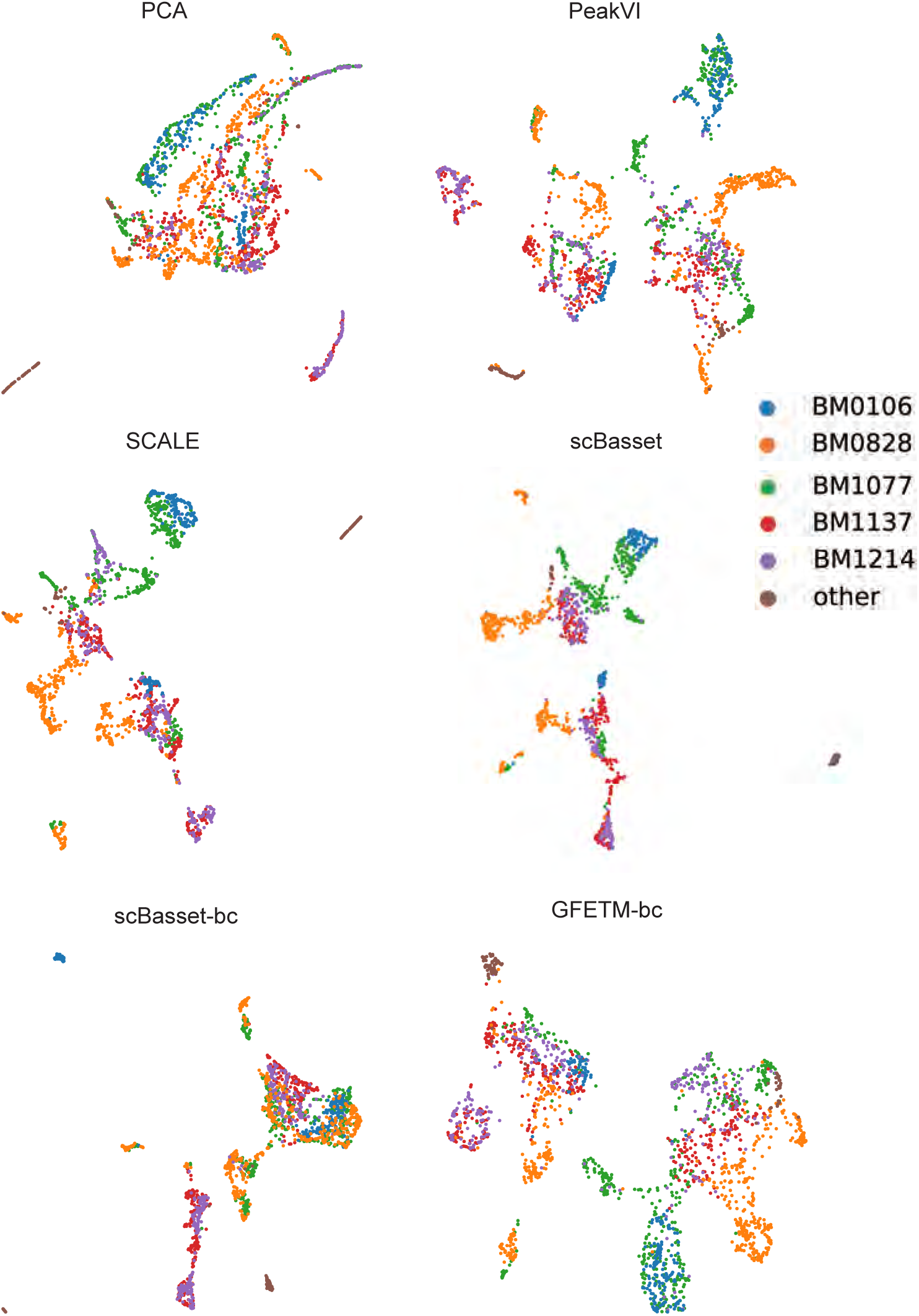
Batch effect correction results of baseline methods on Human HSC Differentiation dataset.

**Supplementary Figure S11:**
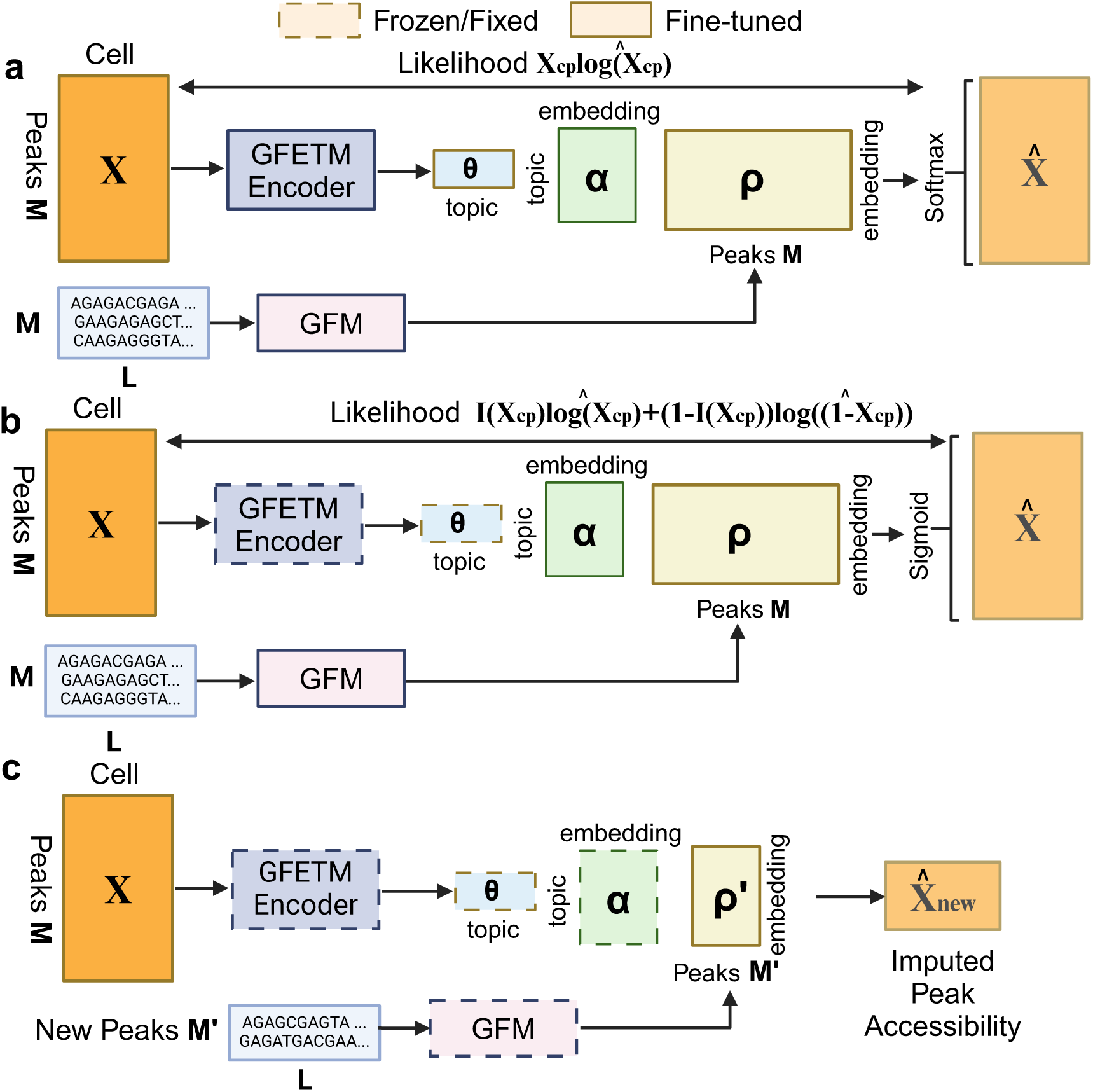
Imputing unseen peaks. The imputation takes 3 stages: a. Standard training of GFETM using the multinomial log likelihood. b. Continued training GFETM to improve imputation accuracy by using the sigmoid output function (σ(θ_c_αρ_p_)) and Bernoulli log likelihood (i.e., binary cross entropy loss). c. Inferring unseen peaks. Given the *M*‘ new sequences, we can infer their likelihood in the N cells by applying the dot product of the cell embedding (θ produced by the trained encoder), topic embedding (α as learned parameters), and peak sequence embedding (ρ‘ produced by the trained GFM).

**Supplementary Figure S12:**
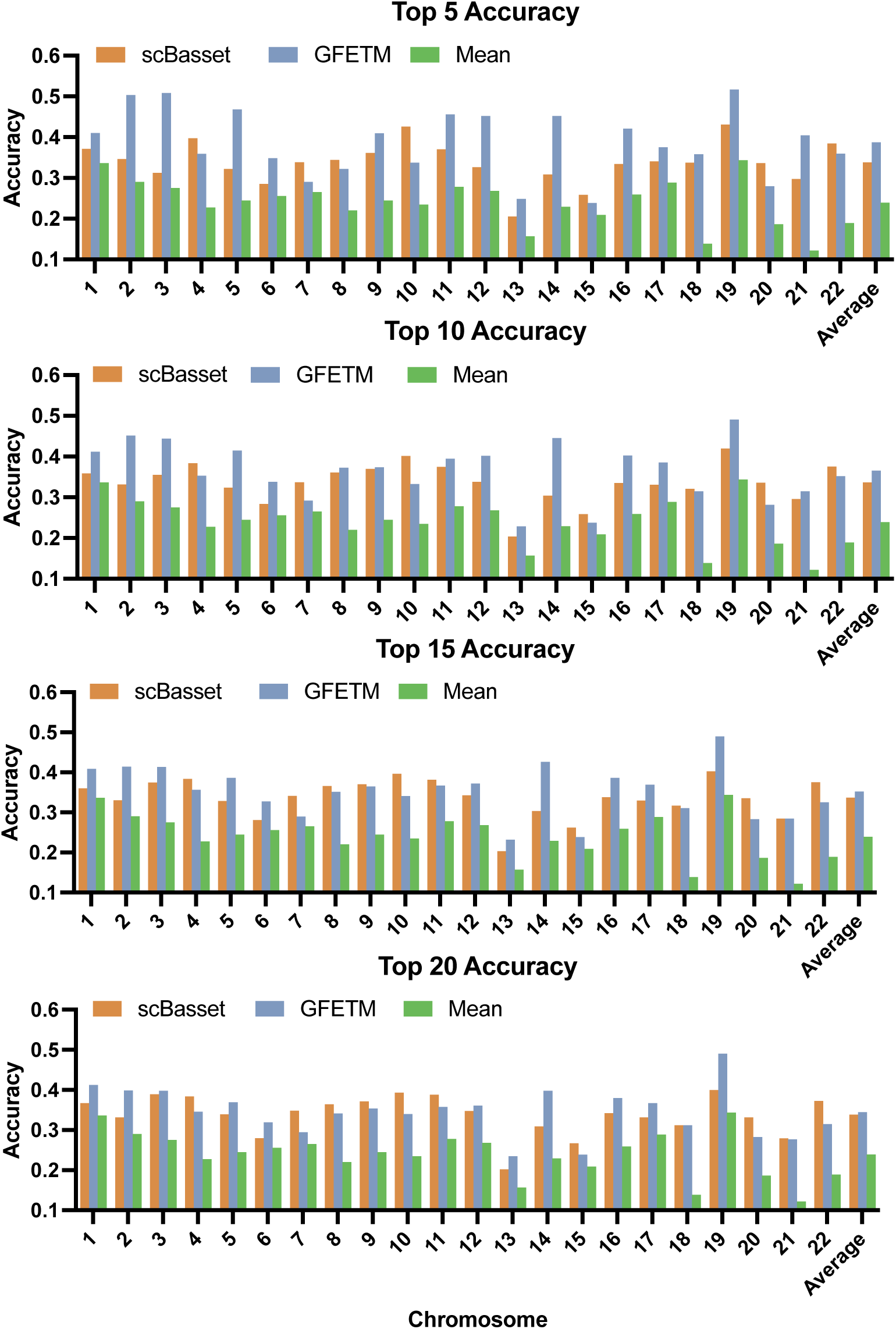
Top K imputation accuracy of unseen peaks. We evaluated scBasset, GFETM, and mean prediction in terms of top K = 5, 10, 15 and 20 accuracy on the 22 chromosomes based on leave-one-chromosome-out cross validation using the Buenrostro2018 dataset.

**Supplementary Figure S13:**
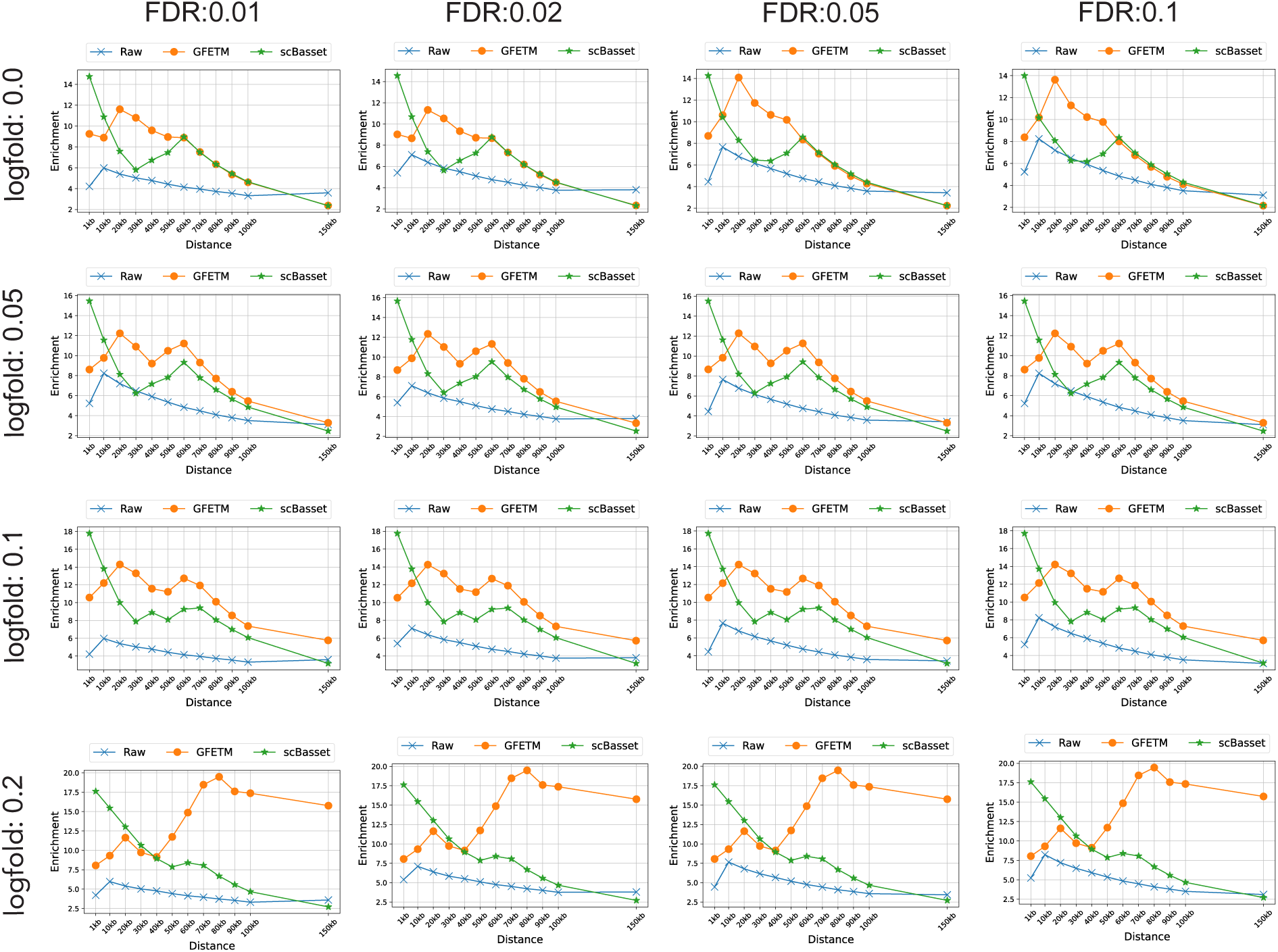
Effects of different hyperparameters (genomic distance, FDR, log-fold change) on the marker gene enrichment score for human HSC cells.

**Supplementary Figure S14:**
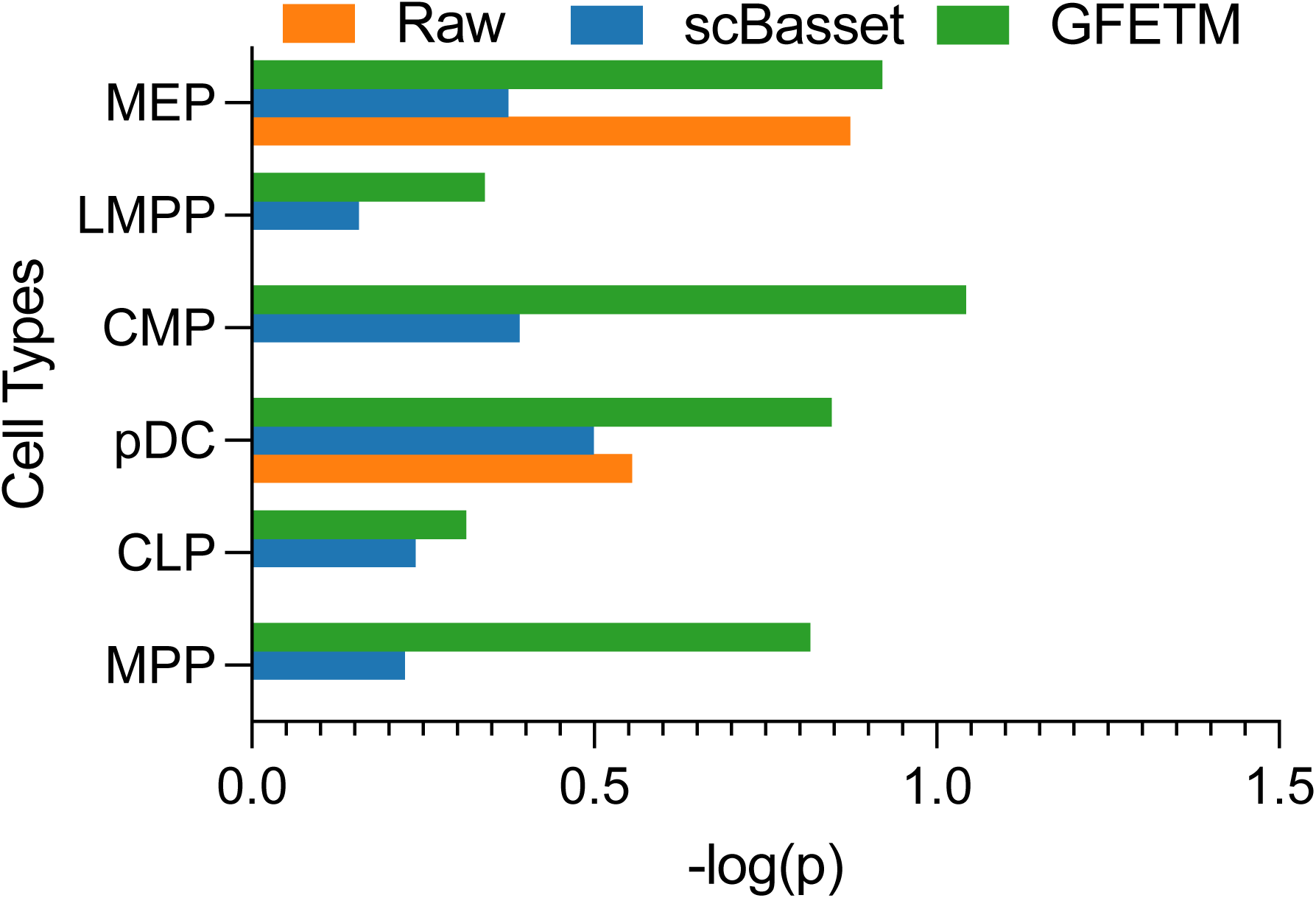
Analysis of the denoised matrix R from GFETM. The enrichment related to marker genes for different cell types are computed from the raw and denoised scATAC matrix. (FDR=0.1, log-fold changes=0, genomic distance=20kb)

**Supplementary Figure S15:**
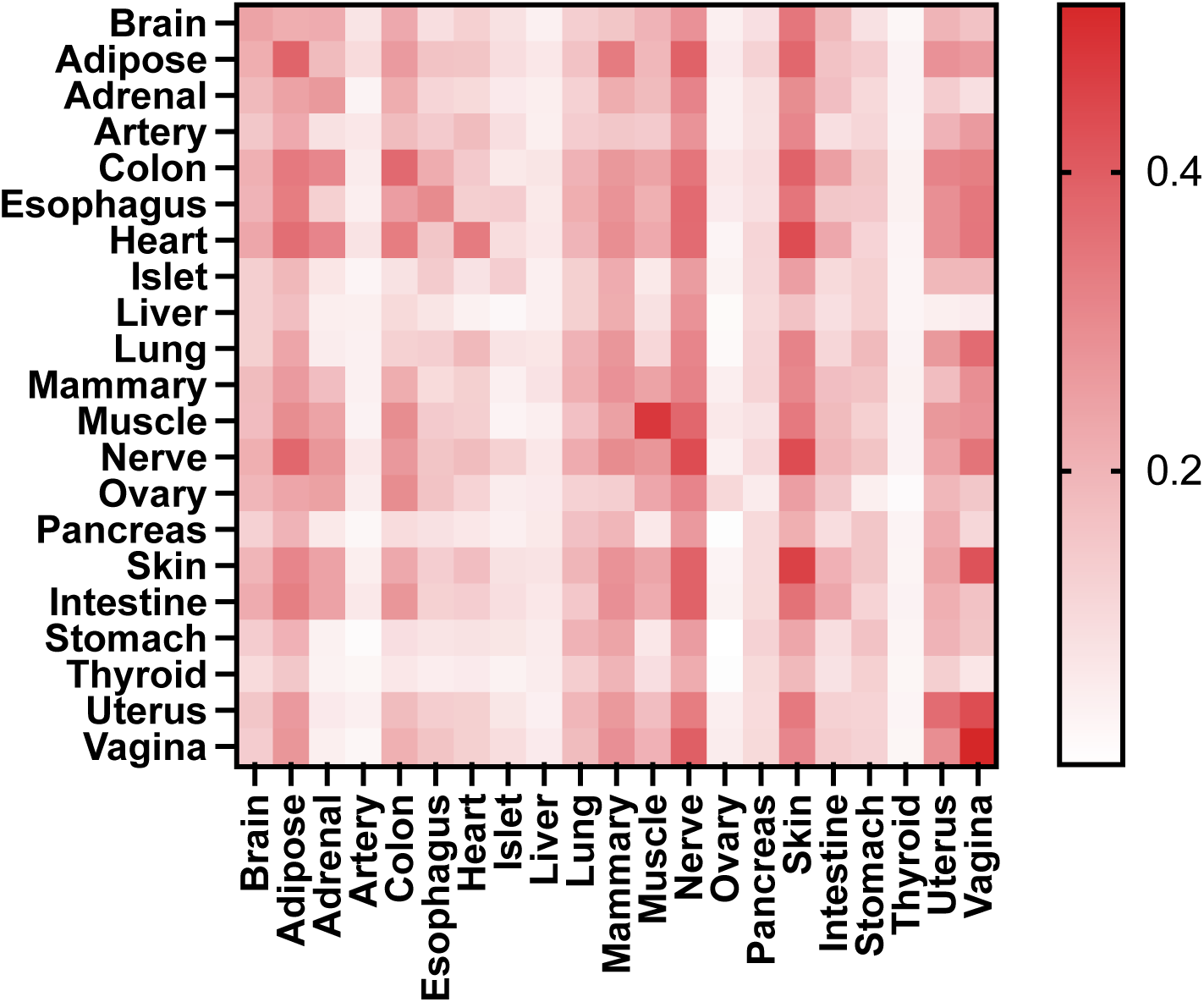
Zero-shot transfer learning on Human tissues

**Supplementary Figure S16:**
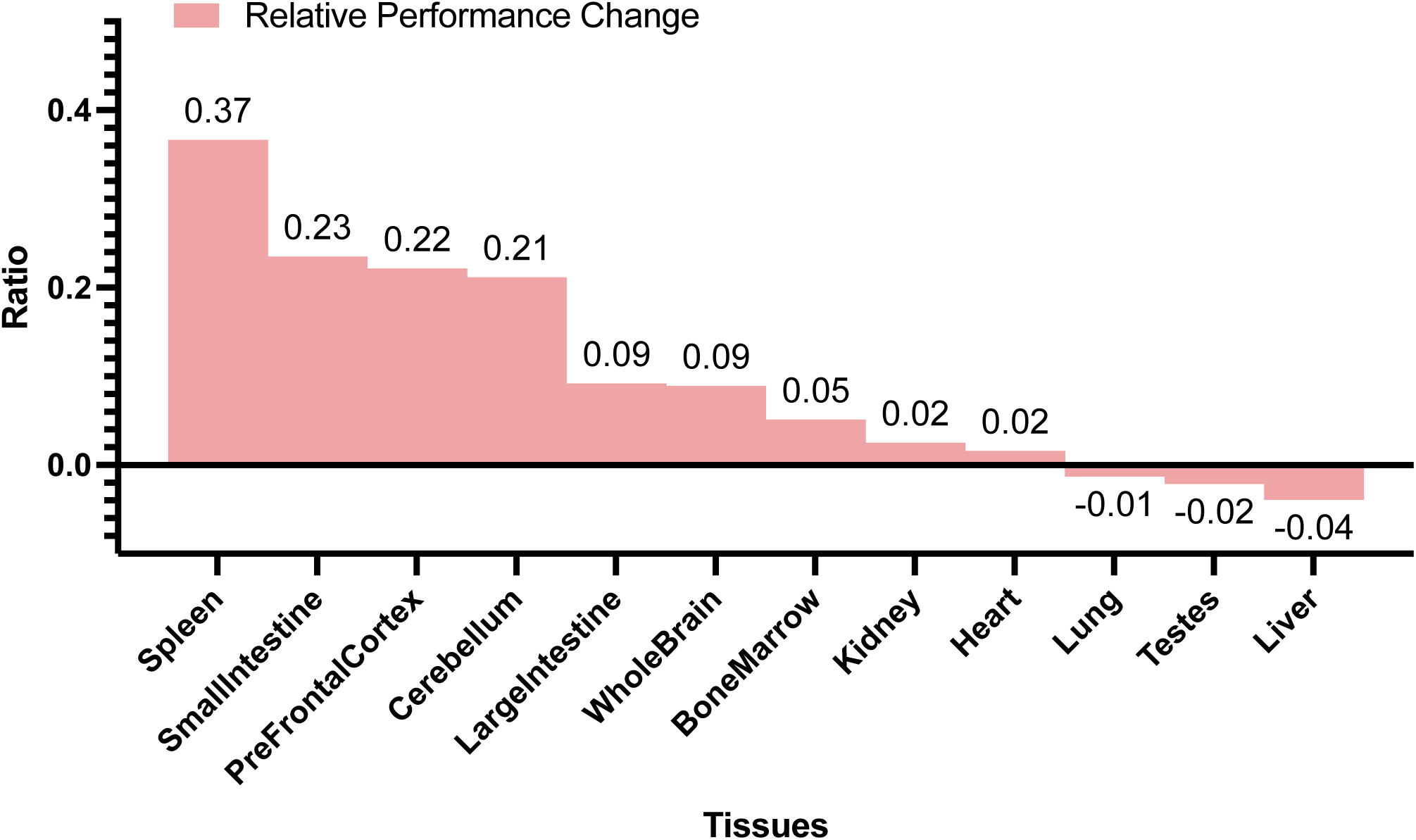
Performance of Transfer+Train on mouse adult tissues.

**Supplementary Figure S17:**
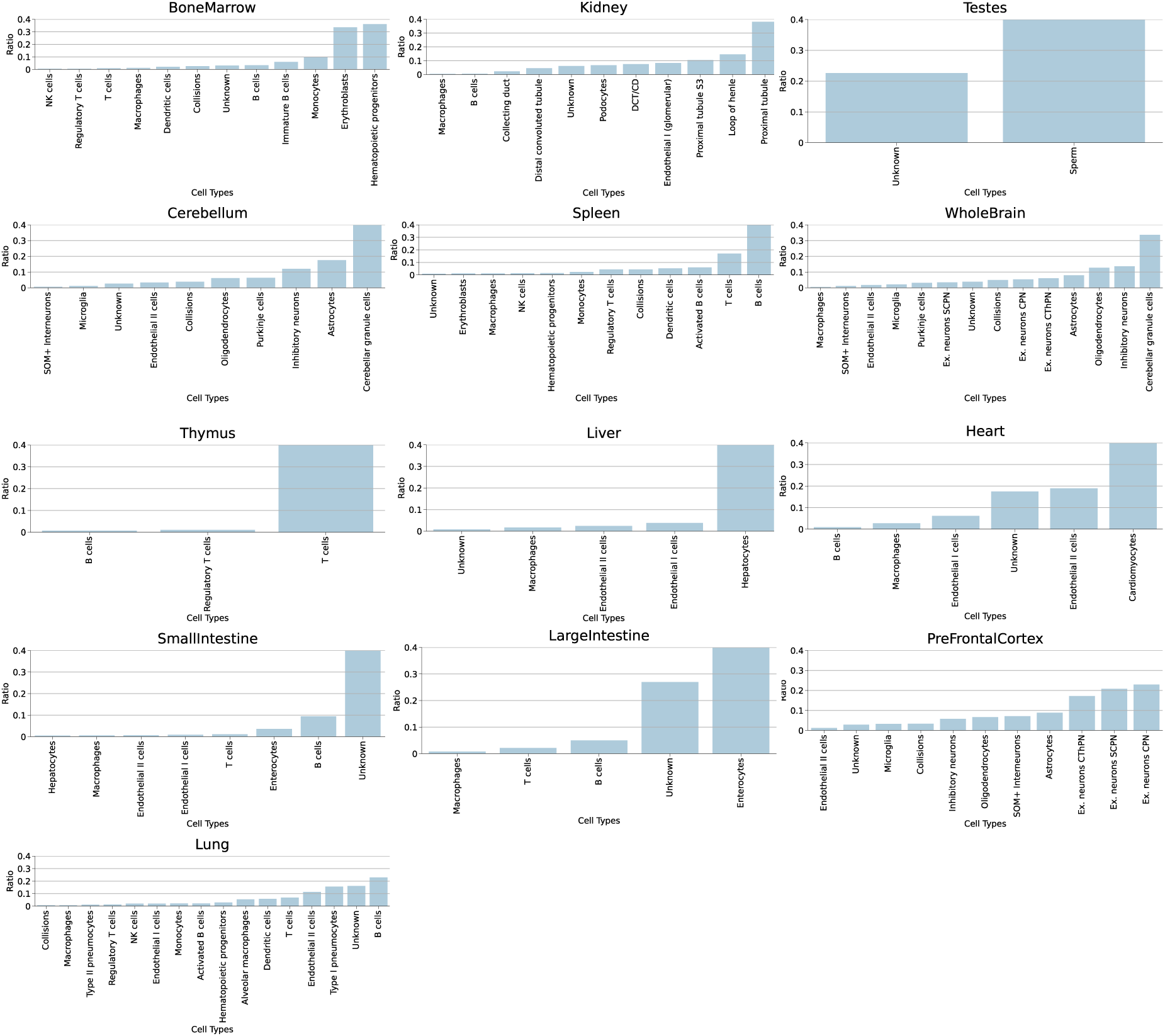
Cell-type proportions of the Cusanovich-mouse dataset.

**Supplementary Figure S18:**
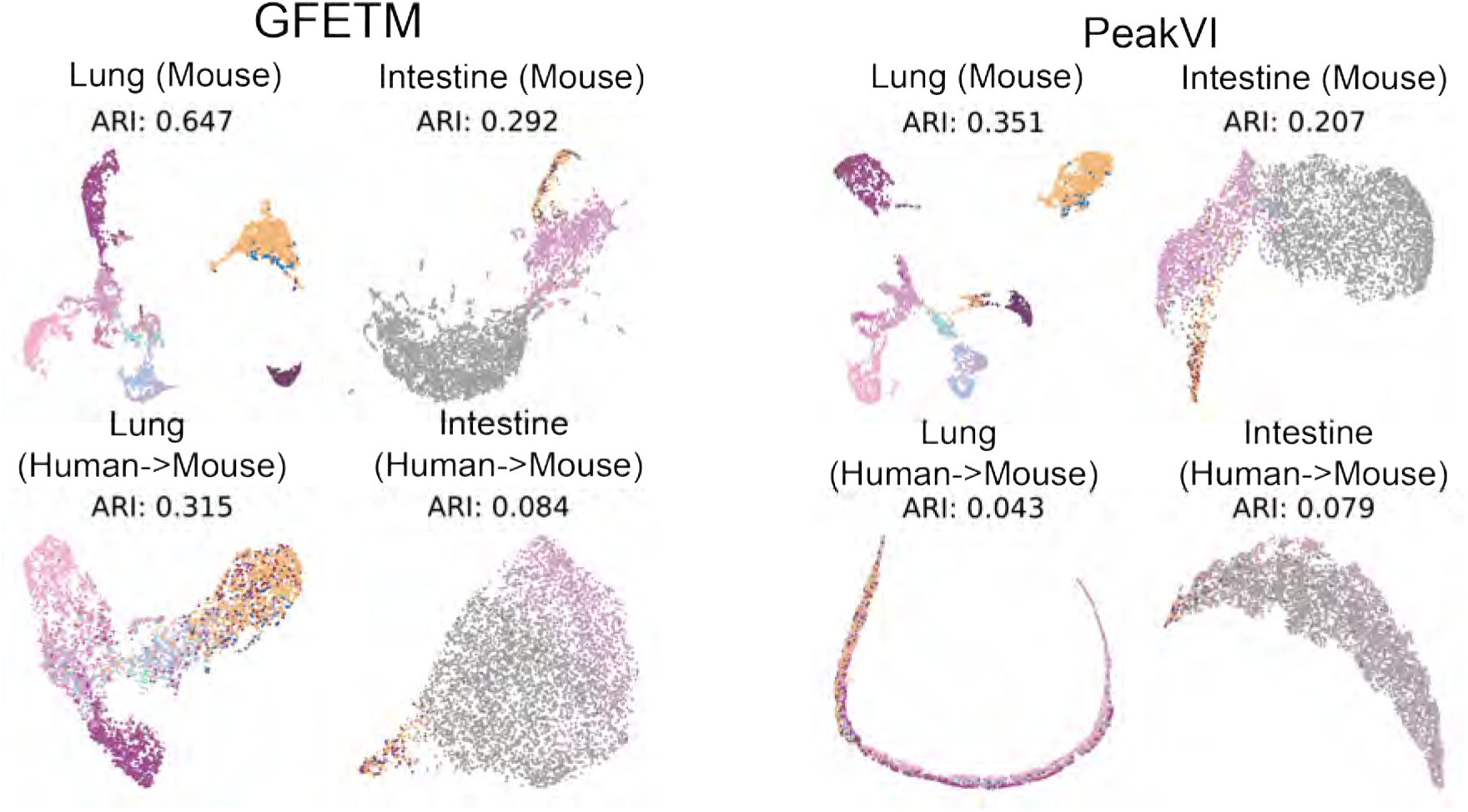
Cross-species transfer between Cusanovich-mouse tissues and Catlas-human tissues for Lung and Intestine

**Supplementary Figure S19:**
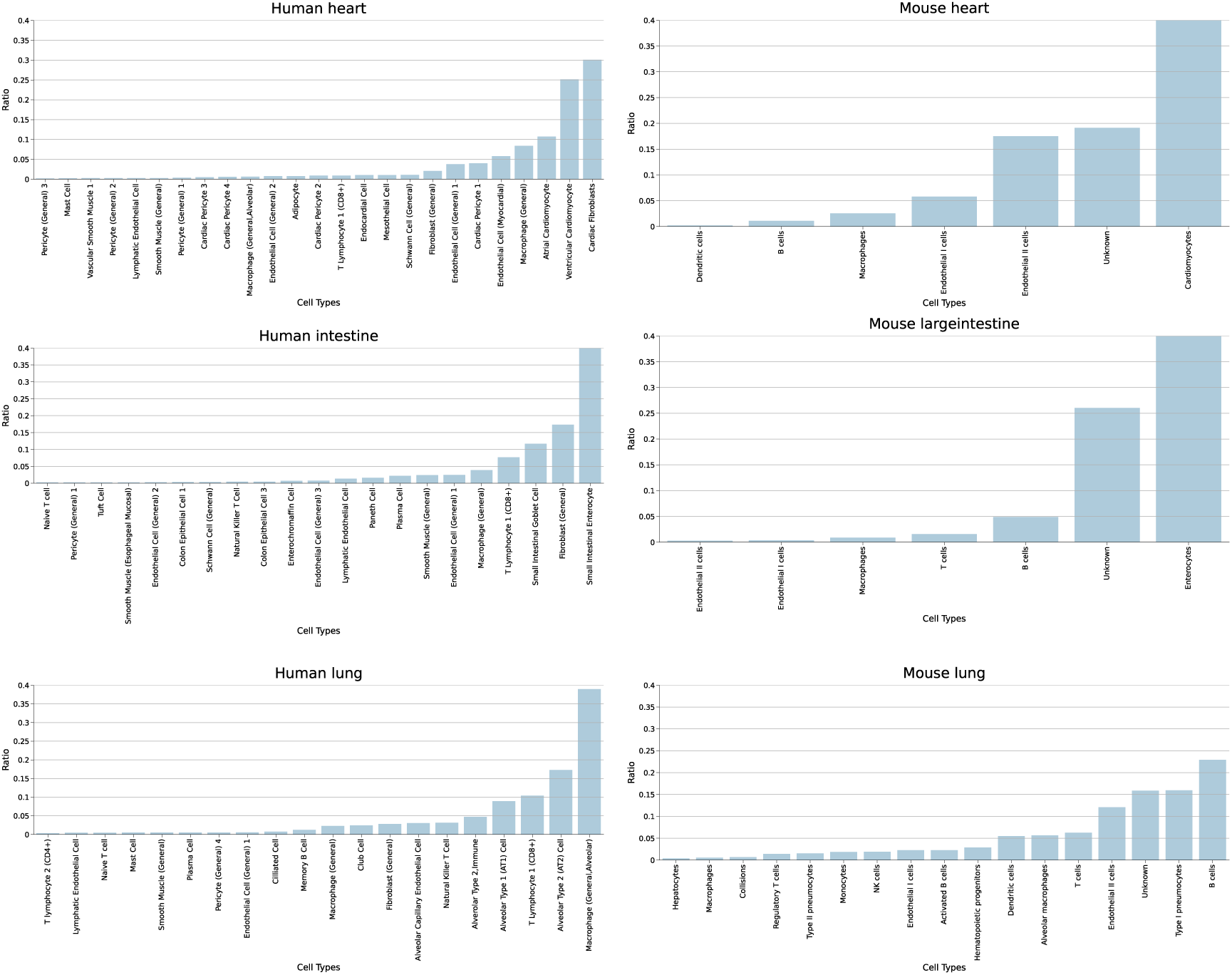
Cell-type proportion of Cusanovich-mouse tissues and Catlas-human tissues included in this study for cross-species transfer learning.

**Supplementary Figure S20:**
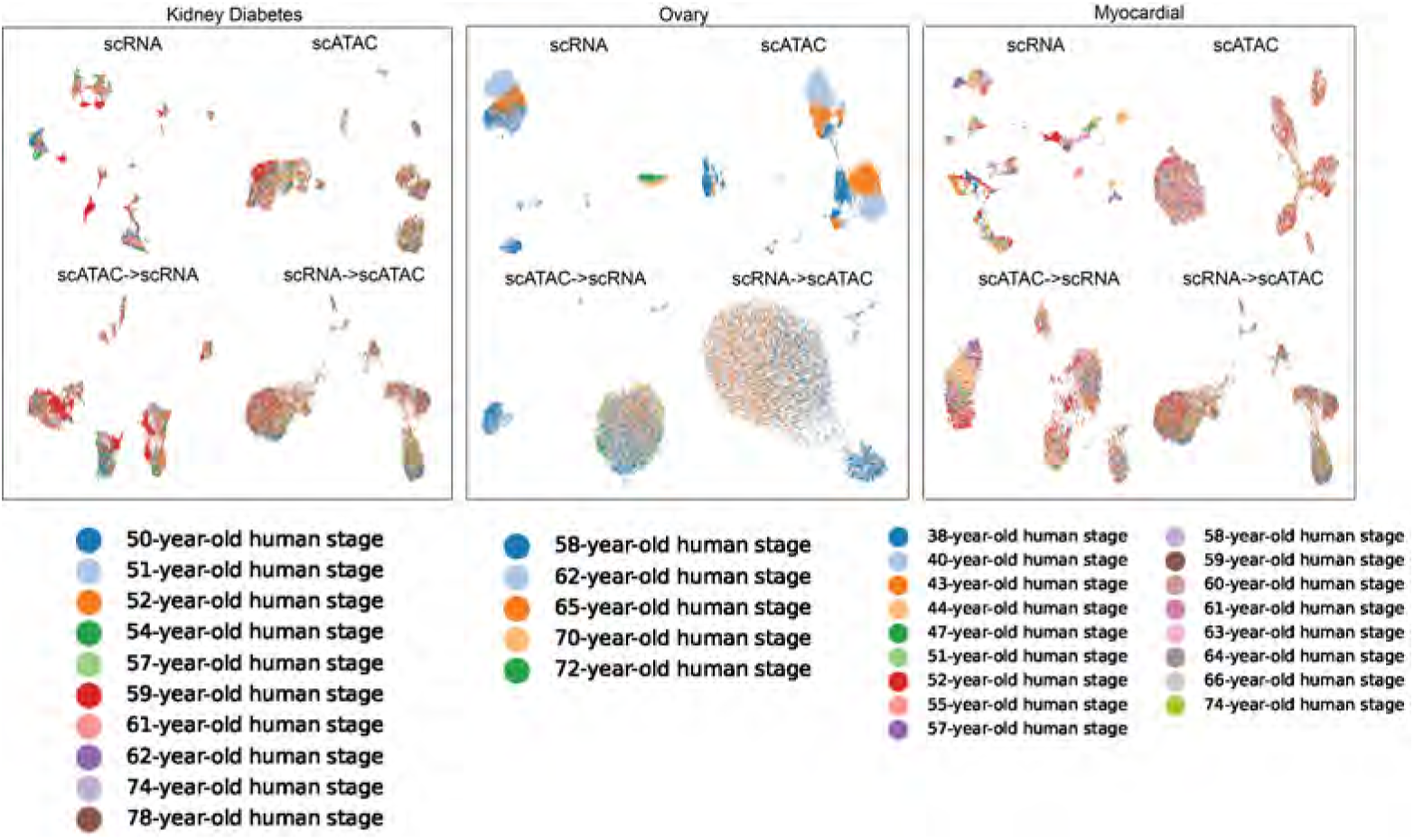
Zero-shot cross-omic transfer learning. Cells were colored by age group.

**Supplementary Figure S21:**
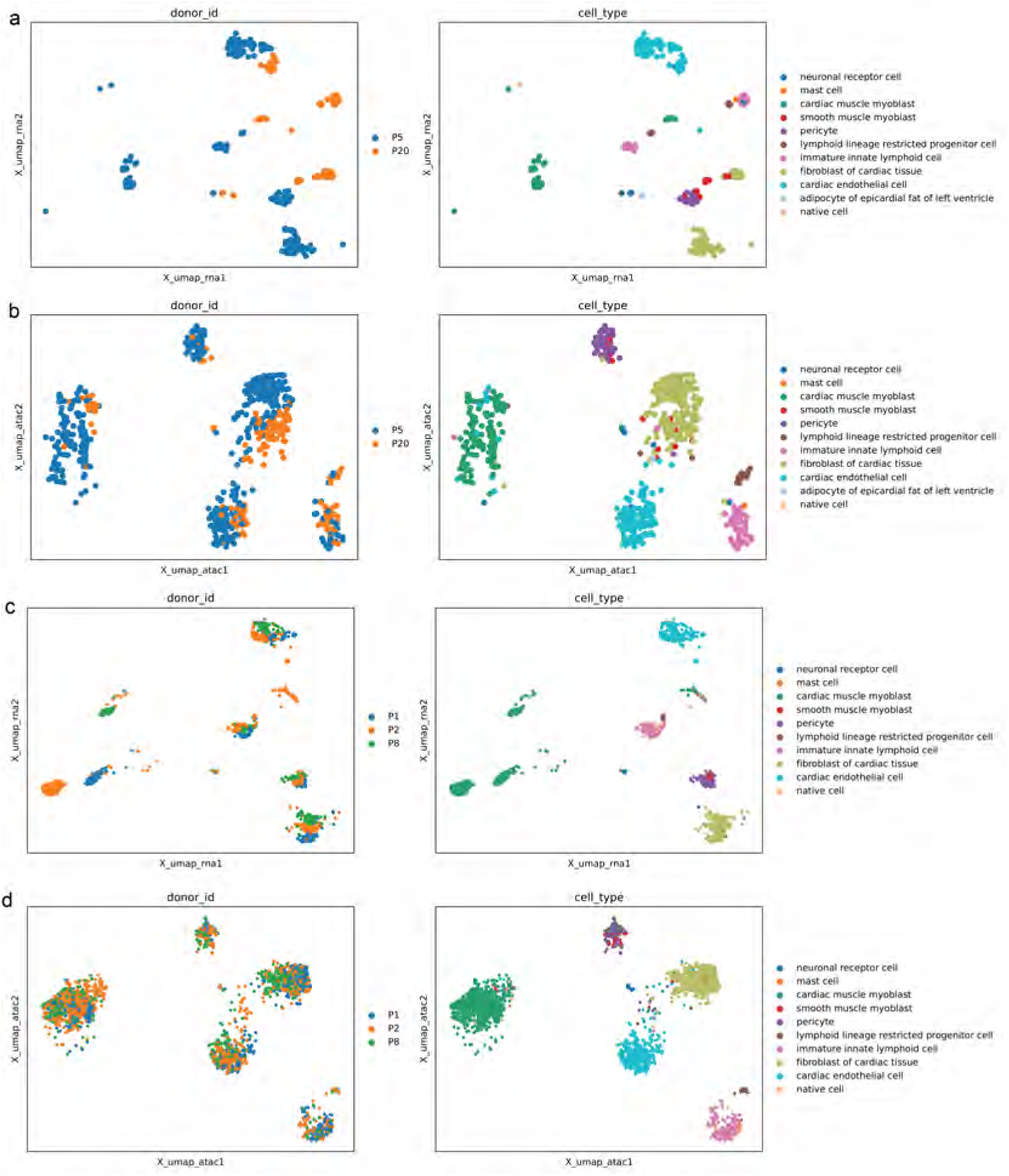
Cross-omic transfer learning on Myocardial dataset. The cell clusters in the left column were colored by donors and the same cell clusters in the right column were colored by cell types. **a.** UMAP visualization of topic embedding from the training scRNA-seq on cells derived from two donors at 63 years of age. We trained GFETM on scRNA-seq data for the myocardial cells from 2 subjects of 63 year old and visualized the cell embedding via UMAP. **b.** Cross-omic transfer learning from scATAC-seq to scRNA-seq on the cells from the same two donors as in panel a. We trained GFETM on the scATAC-seq data of the same subjects as in panel a and applied the trained model to project cells measured by scRNA-seq onto the same topic embedding, which are then visualized via UMAP. **c.** UMAP visualization of topic embedding from the training scRNA-seq dataset on cells derived from 3 donors at 44 years of age. Same as in panel a, we trained GFETM and visualized the embedding on scRNA-seq data. **d.** Cross-omic transfer from scATAC-seq to scRNA-seq on the same cells from the same 3 subjects as in panel c. We trained GFETM on the scATAC-seq data of the same subjects as in panel d and applied the trained model to project cells measured by scRNA-seq onto the same topic embedding, which are then visualized via UMAP.

**Supplementary Figure S22:**
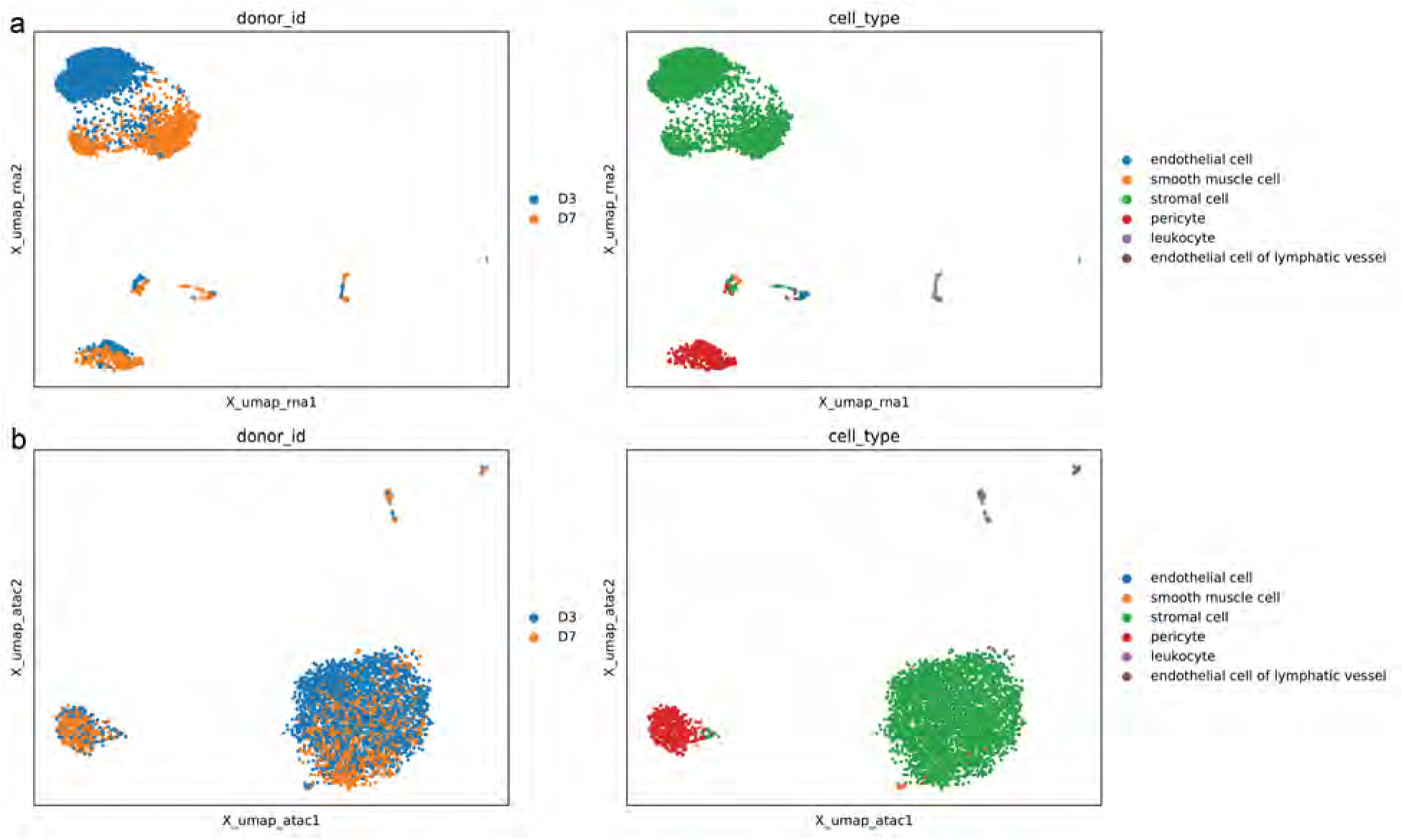
Cross-omic transfer learning on Ovary dataset. The cell clusters in the left column were colored by donors and the same cell clusters in the right column were colored by cell types. **a.** UMAP visualization of topic embedding from the training scRNA-seq on cells derived from two donors at the 62 years of age. We trained GFETM on scRNA-seq data for the ovary cells from two donors and visualized the cell embedding via UMAP. **b.** Cross-omic transfer learning from scATAC-seq to scRNA-seq on the cells from the same two donors as in panel a. We trained GFETM on the scATAC-seq data of the same subjects as in panel a and applied the trained model to project cells measured by scRNA-seq onto the same topic embedding, which are then visualized via UMAP.

**Supplementary Figure S23:**
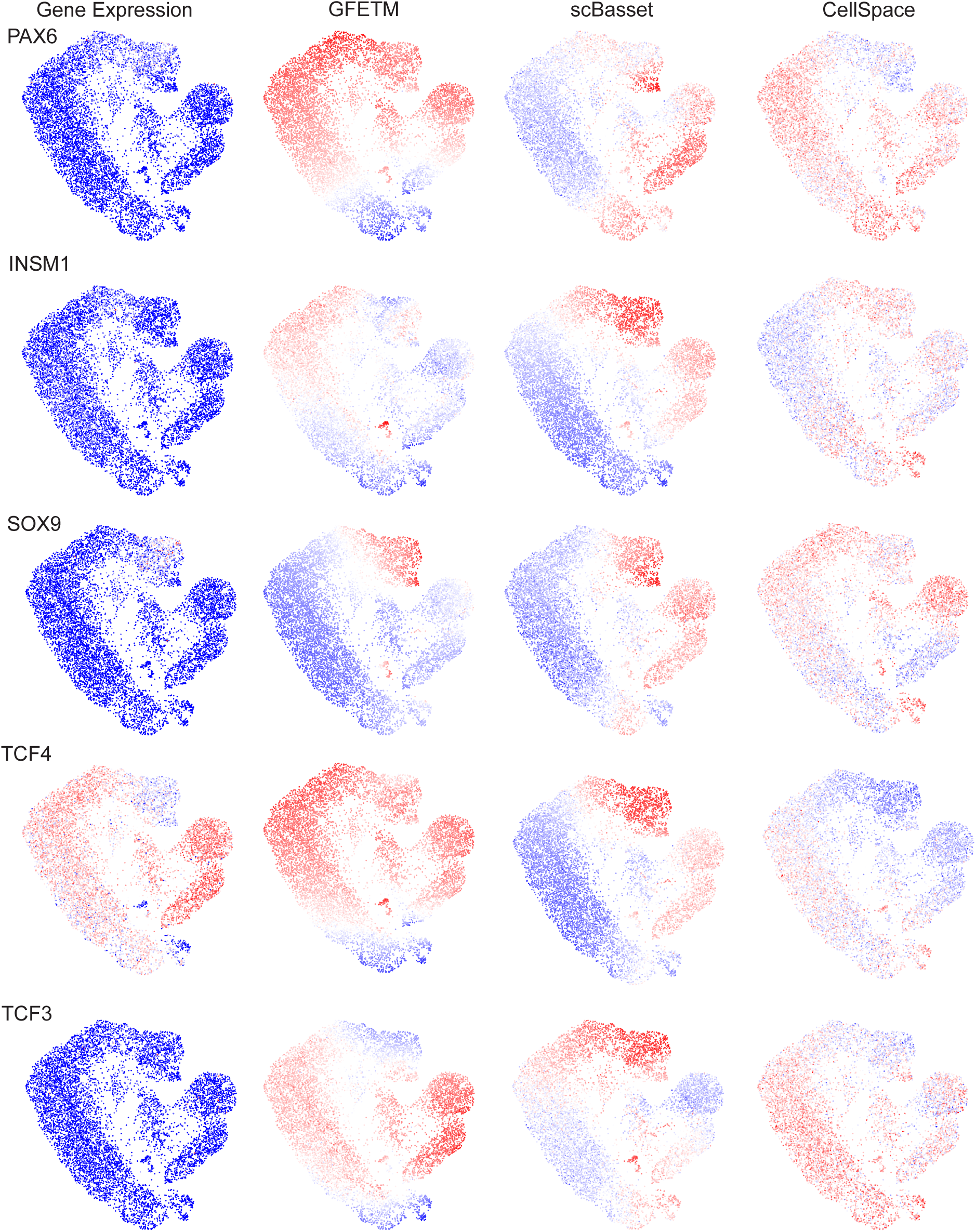
UMAP Visualization of the motif scoring result for PAX6, INSM1, SOX9, TCF4 and TCF3 TFs.

**Supplementary Figure S24:**
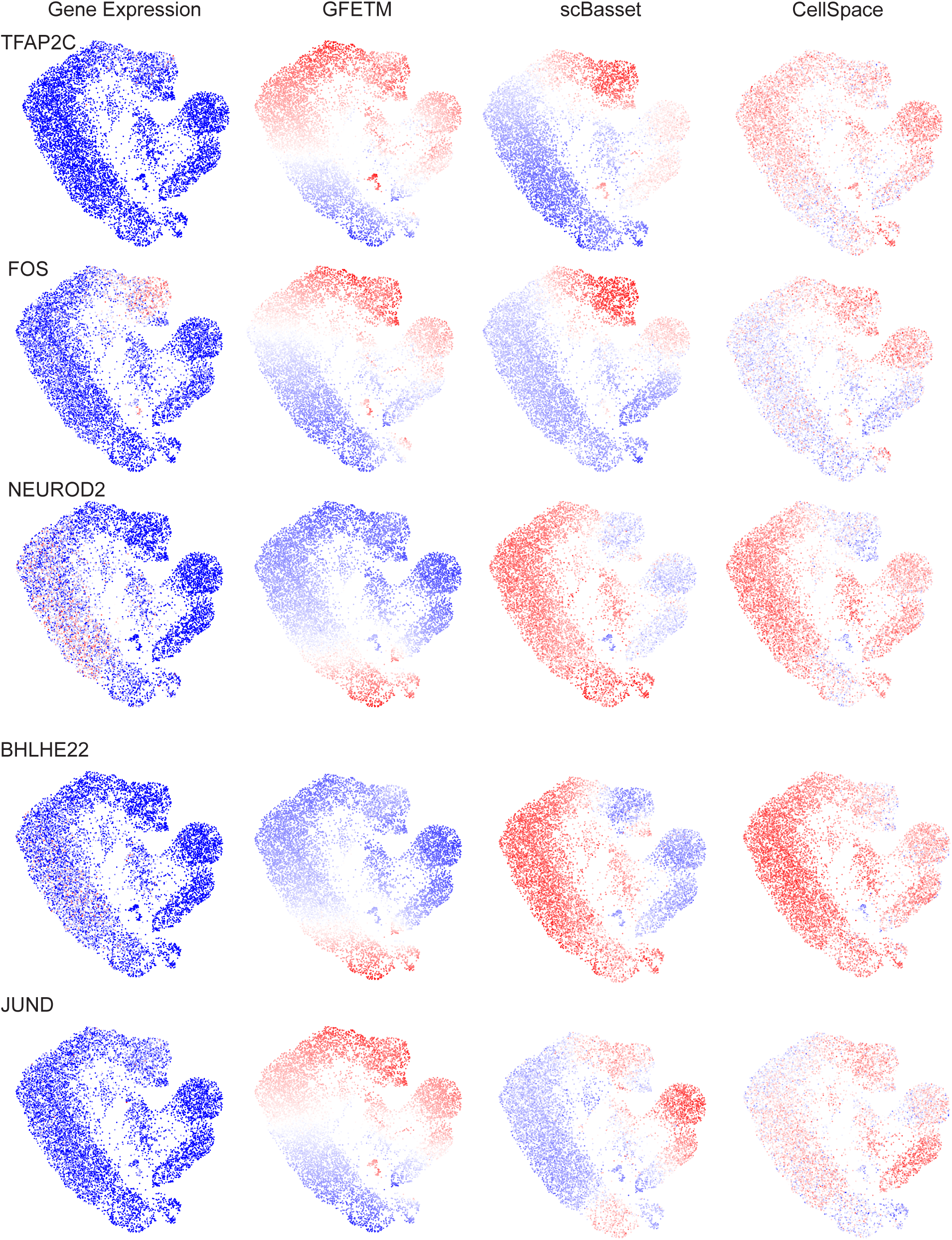
UMAP Visualization of the motif scoring result for TFAP2C, FOS, NEUROD2, BHLHE22 and JUND TFs.

**Supplementary Figure S25:**
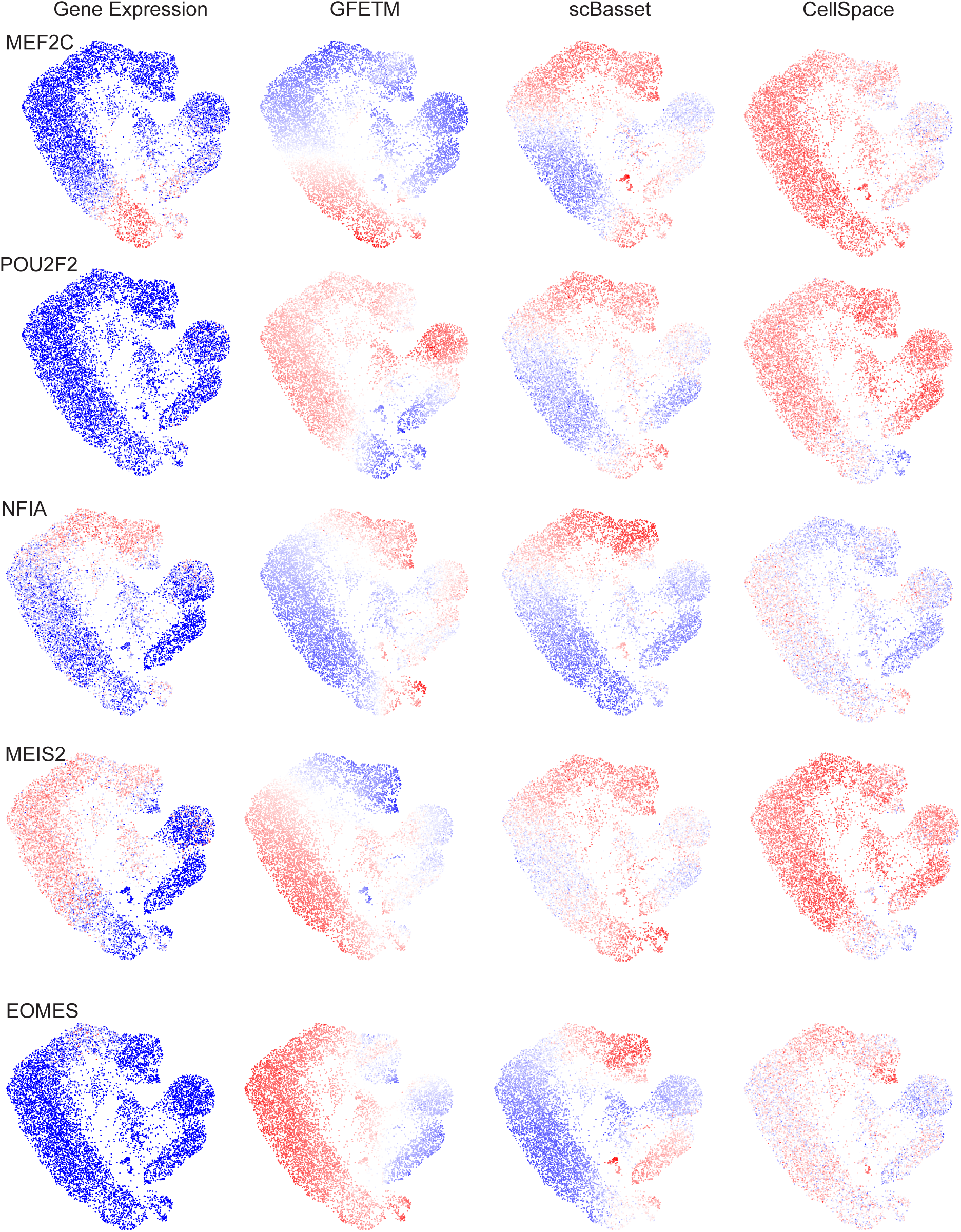
UMAP Visualization of the motif scoring result for MEF2C, POU2F2, NFIA, MEIS2 and EOMES TFs.

**Supplementary Figure S26:**
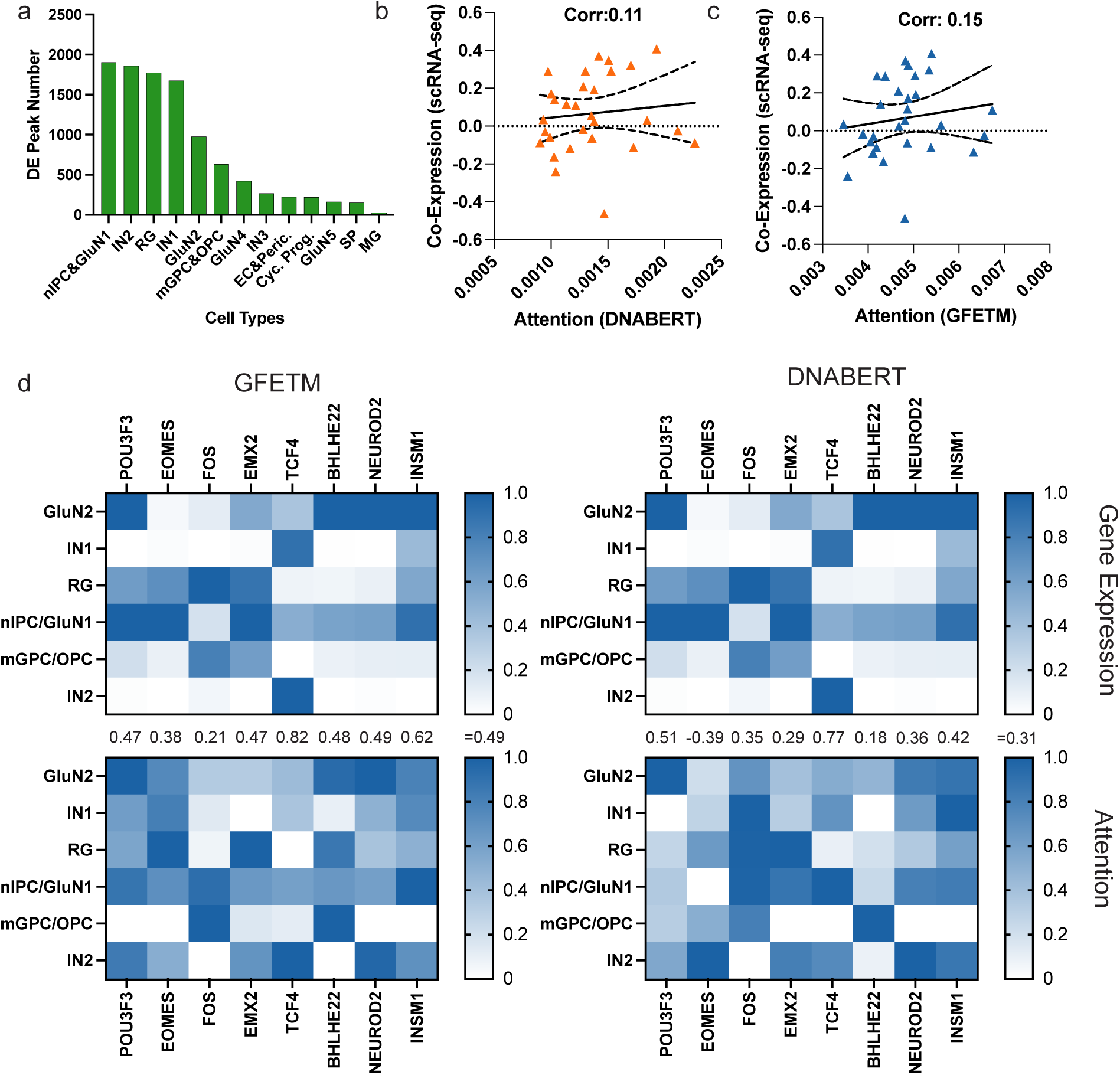
a. Statistics of the DE peak numbers for each cell type in the cortex multiome dataset. b. The correlation between the co-expression weight from scRNA-readout and attention weight from DNABERT. c. The correlation between the co-expression weight from scRNA-seq readout and attention weight from GFETM.

### S1.6 Supplementary Tables

**Supplementary Table S1:** GFMs utilized in this study.

|  | Parameters | Embedding dim | Pre-train Data |
| --- | --- | --- | --- |
| DNABERT-kmer-3 | 86M | 768 | human |
| DNABERT-kmer-4 | 86M | 768 | human |
| DNABERT-kmer-5 | 86M | 768 | human |
| DNABERT-kmer-6 | 89M | 768 | human |
| DNABERT-2 | 117M | 768 | 135 species |
| Nucleotide-Transformers-500m-1000g | 500M | 1280 | human |
| Nucleotide-Transformers-500m-human-ref | 500M | 1280 | human |
| Nucleotide-Transformers-2.5b-1000g | 2.5B | 2560 | human |
| Nucleotide-Transformers-2.5b-multi-speices | 2.5B | 2560 | 850 species |
| hyenadna-tiny-1k-seqlen | 0.4M | 128 | human |
| hyenadna-small-32k-seqlen | 0.4M | 128 | human |
| hyenadna-medium-450k-seqlen | 6.6M | 256 | human |
| hyenadna-large-1m-seqlen | 6.6M | 256 | human |

**Supplementary Table S2:**
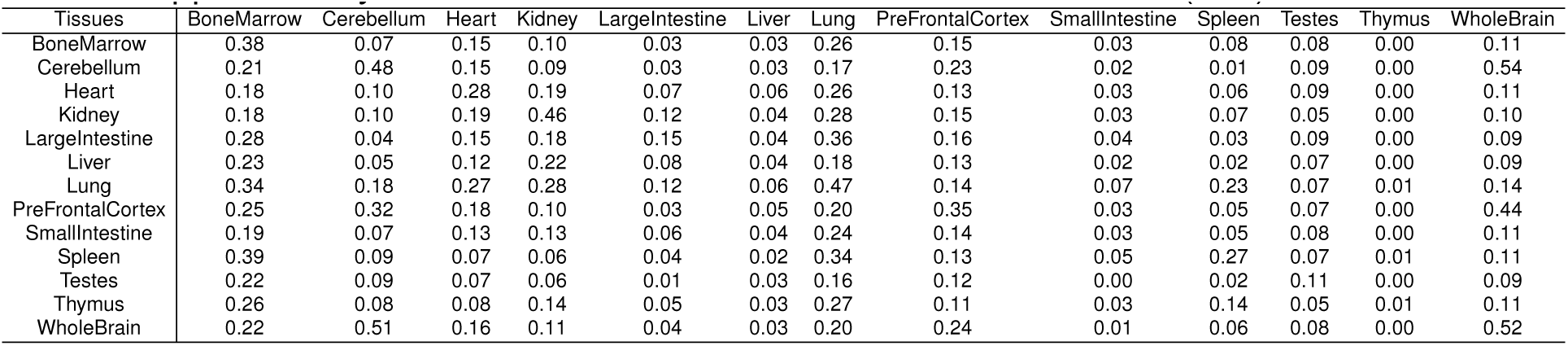
Zero-shot cross-tissue transfer results (ARI) of PeakVI.

**Supplementary Table S3:** Zero-shot cross-tissue transfer results (ARI) of GFETM.

| Tissues | BoneMarrow | Cerebellum | Heart | Kidney | LargeIntestine | Liver | Lung | PreFrontalCortex | SmallIntestine | Spleen | Testes | Thymus | WholeBrain |
| --- | --- | --- | --- | --- | --- | --- | --- | --- | --- | --- | --- | --- | --- |
| BoneMarrow | 0.46 | 0.08 | 0.17 | 0.11 | 0.06 | 0.04 | 0.36 | 0.13 | 0.05 | 0.30 | 0.08 | 0.01 | 0.13 |
| Cerebellum | 0.23 | 0.64 | 0.15 | 0.11 | 0.02 | 0.04 | 0.17 | 0.22 | 0.02 | 0.04 | 0.06 | 0.00 | 0.60 |
| Heart | 0.30 | 0.10 | 0.34 | 0.22 | 0.08 | 0.06 | 0.41 | 0.17 | 0.04 | 0.14 | 0.08 | 0.01 | 0.12 |
| Kidney | 0.30 | 0.10 | 0.28 | 0.46 | 0.14 | 0.05 | 0.40 | 0.16 | 0.05 | 0.11 | 0.06 | 0.00 | 0.16 |
| LargeIntestine | 0.33 | 0.09 | 0.23 | 0.21 | 0.20 | 0.05 | 0.42 | 0.14 | 0.08 | 0.18 | 0.07 | 0.00 | 0.11 |
| Liver | 0.27 | 0.09 | 0.21 | 0.25 | 0.11 | 0.06 | 0.30 | 0.11 | 0.02 | 0.10 | 0.05 | 0.00 | 0.11 |
| Lung | 0.37 | 0.14 | 0.28 | 0.25 | 0.16 | 0.06 | 0.58 | 0.16 | 0.06 | 0.31 | 0.10 | 0.02 | 0.14 |
| PreFrontalCortex | 0.24 | 0.37 | 0.21 | 0.13 | 0.04 | 0.04 | 0.25 | 0.36 | 0.03 | 0.11 | 0.09 | 0.01 | 0.43 |
| SmallIntestine | 0.24 | 0.07 | 0.15 | 0.14 | 0.04 | 0.04 | 0.30 | 0.13 | 0.06 | 0.11 | 0.08 | 0.00 | 0.10 |
| Spleen | 0.36 | 0.07 | 0.09 | 0.09 | 0.05 | 0.03 | 0.31 | 0.12 | 0.06 | 0.33 | 0.07 | 0.01 | 0.08 |
| Testes | 0.21 | 0.04 | 0.06 | 0.06 | 0.01 | 0.03 | 0.07 | 0.11 | 0.02 | 0.03 | 0.13 | 0.00 | 0.07 |
| Thymus | 0.27 | 0.06 | 0.09 | 0.07 | 0.05 | 0.03 | 0.30 | 0.12 | 0.05 | 0.21 | 0.09 | 0.01 | 0.08 |
| WholeBrain | 0.24 | 0.55 | 0.20 | 0.12 | 0.04 | 0.04 | 0.22 | 0.27 | 0.04 | 0.08 | 0.10 | 0.01 | 0.66 |

**Supplementary Table S4:** Zero-shot cross-omic transfer results (ARI) of PeakVI.

| Dataset | rna | atac | rna->atac | atac->rna |
| --- | --- | --- | --- | --- |
| Myocardial | 0.39 | 0.47 | 0.39 | 0.49 |
| Ovary | 0.14 | 0.09 | 0.08 | 0.11 |
| Kidney Diabetic | 0.48 | 0.49 | 0.57 | 0.51 |

**Supplementary Table S5:** ARI clustering metrics of GFETM and two baseline methods on the Cusanovich-mouse dataset (20000 cells). 10,20,30,40 indicate the number of clusters in the louvain clustering algorithm. 1024,2048,…,65536 indicate the number of selected highly variable peaks.

| GFETM | 10 | 20 | 30 | 40 | PeakVI | 10 | 20 | 30 | 40 | scBasset | 10 | 20 | 30 | 40 |
| --- | --- | --- | --- | --- | --- | --- | --- | --- | --- | --- | --- | --- | --- | --- |
| 1024 | 0.11 | 0.09 | 0.09 | 0.08 | 1024 | 0.12 | 0.13 | 0.11 | 0.10 | 1024 | 0.12 | 0.14 | 0.14 | 0.14 |
| 2048 | 0.18 | 0.16 | 0.15 | 0.15 | 2048 | 0.19 | 0.21 | 0.21 | 0.17 | 2048 | 0.23 | 0.27 | 0.29 | 0.25 |
| 4096 | 0.26 | 0.24 | 0.23 | 0.23 | 4096 | 0.28 | 0.34 | 0.31 | 0.26 | 4096 | 0.30 | 0.43 | 0.39 | 0.40 |
| 8192 | 0.37 | 0.45 | 0.42 | 0.42 | 8192 | 0.31 | 0.52 | 0.46 | 0.41 | 8192 | 0.35 | 0.52 | 0.56 | 0.56 |
| 16384 | 0.45 | 0.66 | 0.63 | 0.59 | 16384 | 0.37 | 0.62 | 0.60 | 0.53 | 16384 | 0.36 | 0.60 | 0.60 | 0.60 |
| 32768 | 0.45 | 0.68 | 0.63 | 0.61 | 32768 | 0.38 | 0.65 | 0.59 | 0.53 | 32768 | 0.411 | 0.64 | 0.60 | 0.56 |
| 65536 | 0.46 | **0.75** | 0.67 | 0.59 | 65536 | 0.42 | **0.69** | 0.64 | 0.54 | 65536 | 0.355 | 0.66 | **0.67** | 0.55 |

**Supplementary Table S6:** ARI clustering metrics of GFETM and two baseline methods on the Cusanovich-mouse dataset (60000 cells). 10,20,30,40 indicate the number of clusters in the louvain clustering algorithm. 1024,2048,…,65536 indicate the number of selected highly variable peaks.

| GFETM | 10 | 20 | 30 | 40 | PeakVI | 10 | 20 | 30 | 40 | scBasset | 10 | 20 | 30 | 40 |
| --- | --- | --- | --- | --- | --- | --- | --- | --- | --- | --- | --- | --- | --- | --- |
| 1024 | 0.14 | 0.12 | 0.11 | 0.13 | 1024 | 0.14 | 0.17 | 0.15 | 0.13 | 1024 | 0.17 | 0.15 | 0.14 | 0.14 |
| 2048 | 0.23 | 0.19 | 0.18 | 0.18 | 2048 | 0.13 | 0.26 | 0.24 | 0.20 | 2048 | 0.26 | 0.32 | 0.33 | 0.33 |
| 4096 | 0.30 | 0.28 | 0.27 | 0.26 | 4096 | 0.26 | 0.43 | 0.39 | 0.33 | 4096 | 0.33 | 0.44 | 0.39 | 0.39 |
| 8192 | 0.38 | 0.55 | 0.47 | 0.44 | 8192 | 0.33 | 0.53 | 0.48 | 0.42 | 8192 | 0.31 | 0.54 | 0.53 | 0.53 |
| 16384 | 0.45 | 0.68 | 0.63 | 0.58 | 16384 | 0.37 | 0.62 | 0.61 | 0.57 | 16384 | 0.37 | 0.64 | 0.59 | 0.55 |
| 32768 | 0.43 | 0.70 | 0.69 | 0.64 | 32768 | 0.43 | 0.65 | 0.64 | 0.64 | 32768 | 0.37 | 0.64 | 0.68 | 0.53 |
| 65536 | 0.47 | **0.72** | 0.71 | 0.66 | 65536 | 0.40 | **0.71** | 0.69 | 0.64 | 65536 | 0.41 | 0.66 | **0.68** | 0.55 |

## Footnotes

1 https://biopython.org/

## Notes

### Competing Interest Statement

The authors have declared no competing interest.

### Summary of Updates

Added TF motif activities analysis. Improved figure quality and overall writing.

